# Spatially patterned mitochondrial remodeling and metabolic changes during Müller glia-dependent retinal regeneration in zebrafish

**DOI:** 10.64898/2026.11.10.765195

**Authors:** Reza Raeisossadati, Michael A. Robichaux, Jeffrey M. Gross

## Abstract

Müller glia regenerate retinal neurons after injury in zebrafish, but how their mitochondrial organization changes during regenerative activation remains poorly defined. Here, we combined mitochondrial imaging, Müller glial transcriptomics, and whole eyecup metabolomics to investigate the response to selective photoreceptor loss. A Müller glia mitochondrial reporter revealed spatial and temporal mitochondrial remodeling: morphological changes were detectable in inner retinal compartments before sustained increases in mitochondrial volume emerged in processes adjacent to injured photoreceptors. This response coincided with reduced respiratory chain transcript expression and selective induction of mitochondrial assembly and maintenance genes, while the eyecup underwent temporally patterned metabolic changes. Dichloroacetate (DCA), an inhibitor of pyruvate dehydrogenase kinases (PDK), attenuated outer-retinal Müller glial mitochondrial remodeling. DCA also reshaped the Müller glial transcriptional response to injury, suppressing reprogramming- and cell-cycle-associated programs and altering metabolic and mitochondrial gene expression. DCA treatment also blocked the formation of *tuba1a*:GFP^+^ Müller glia and PCNA expression, indicating that reprogramming and progenitor proliferation requires PDK activity. Together, our findings identify compartment-specific mitochondrial remodeling as an early event in the Müller glial regenerative response and demonstrate that mitochondrial remodeling and regenerative activation are sensitive to perturbation of the PDK–PDH regulatory axis.

**Significance Statement:** Zebrafish Müller glia can generate progenitors that replace injured retinal neurons, but how their mitochondria reorganize during this transition remains poorly understood. We show that localized photoreceptor loss elicits mitochondrial remodeling across Müller glial compartments, with distinct temporal responses that accompany selective mitochondrial gene regulation and tissue-level metabolic changes. Inhibiting pyruvate dehydrogenase kinases attenuates outer-retinal mitochondrial remodeling and markedly reduces reprogramming-associated and proliferative responses. These findings identify spatial mitochondrial organization as a component of regenerative glial activation and establish a framework for investigating how metabolic regulation influences endogenous retinal repair.

## Introduction

Retinal injury triggers Müller glia-dependent neuronal regeneration in zebrafish, whereas the endogenous regenerative capacity of mammalian Müller glia is limited [1, 2]. In zebrafish, Müller glia activate transcriptional programs that enable reprogramming and the production of progenitors that regenerate retinal neurons lost to injury [3]. Comparative studies in zebrafish and mice have identified signaling pathways and transcriptional and epigenetic regulatory networks that promote this transition in zebrafish and constrain it in mammals [1]. Experimental reprogramming can partially overcome these constraints in mouse Müller glia, raising the possibility of harnessing their latent neurogenic potential for retinal repair in mammals [2]. Despite progress in defining the regulatory programs underlying Müller glial-dependent retinal regeneration, how Müller glia reorganize their mitochondria and metabolism during the transition toward a regenerative state remains poorly understood.

There is evidence that mitochondria connect cellular responses to injury with the transition toward a regenerative state. Beyond their roles in energy production, mitochondrial dynamics can influence stem cell self-renewal and fate decisions [4]. Metabolic regulation also influences regenerative proliferation: in the injured zebrafish heart, glycolysis supports cardiomyocyte proliferation, while pharmacological inhibition of pyruvate dehydrogenase kinase (PDK) activity reduces this response [5]. In the retina, stressed cone photoreceptors transfer damaged mitochondria to Müller glia, linking neuronal mitochondrial stress to glial handling of damaged organelles [6]. Whether Müller glia remodel their own mitochondrial population during the injury and regenerative response remains poorly defined, however.

Müller glia span all layers of the retina, placing their endfeet, somata, and processes in contact with distinct neuronal populations. These neighboring populations can differ metabolically: rod photoreceptors require both glycolysis and oxidative phosphorylation to maintain ATP levels, whereas ganglion and amacrine cells rely predominantly on oxidative phosphorylation [7]. This heterogeneity motivates asking whether mitochondrial responses within Müller glia differ across retinal compartments. Indeed, while whole-cell transcriptional analyses can identify injury-responsive mitochondrial programs, they cannot resolve how mitochondrial organization changes within these distinct compartments. Studies of neuronal regeneration illustrate the importance of this spatial dimension. In adult zebrafish retinal ganglion cells, optic nerve injury induces different changes in mitochondrial morphology and abundance across dendritic, somatic, and axonal compartments [8]. Moreover, enhancing mitochondrial transport and positioning in injured mammalian neurons improves axonal regeneration and alleviates local energy deficits, demonstrating that mitochondrial positioning can influence regenerative growth [9].

Here, we developed a zebrafish *gfap*:mito-GFP knock-in reporter to image Müller glial mitochondrial morphology across retinal compartments following selective ablation of ultraviolet (UV)-sensitive cone photoreceptors. We combined compartment-resolved NSPARC super-resolution imaging with RNA sequencing of FACS-isolated Müller glia and whole eyecup metabolomics to examine mitochondrial remodeling during the injury response. We further perturbed the mitochondrial pyruvate dehydrogenase kinase– pyruvate dehydrogenase (PDK–PDH) axis using dichloroacetate (DCA). Our findings reveal spatially and temporally patterned mitochondrial responses within Müller glia and show that DCA attenuates outer-retinal mitochondrial remodeling while reducing reprogramming-associated gene expression and proliferative activation in Müller glia. Together, these results identify mitochondrial remodeling as an early component of the Müller glial regenerative response and demonstrate its sensitivity to perturbation of metabolic regulation.

## Results

### Photoreceptor loss induces spatially and temporally patterned mitochondrial remodeling in Müller glia

Because Müller glia span all layers of the retina, photoreceptor loss could elicit different mitochondrial responses in their somata and processes. To resolve potential responses across retinal compartments, we generated a transgenic reporter line, *gfap*^Tg(MITO-GFP)^ (hereafter mito-GFP), that labels mitochondria in Müller glia (SI Fig. 1). We crossed mito-GFP fish with *Tg*(*sws1*-nsfb-mCherry) fish [10] to enable selective ablation of UV-sensitive cone photoreceptors. In cells expressing nitroreductase (NfsB), metronidazole (MTZ) is converted into a cytotoxic metabolite, resulting in ablation of the NfsB-expressing cells [11]. Adult fish were exposed to 10 mM MTZ for 16h, and Müller glial mitochondrial morphology was imaged and quantified at 24, 48, and 60h after treatment (hours post treatment; hpt) (Fig. 1A,B). We quantified mitochondrial morphology separately within the ganglion cell layer (GCL), inner plexiform layer (IPL), inner nuclear layer (INL), outer plexiform layer (OPL), and outer nuclear layer (ONL) (Fig. 1C-F; raw images SI Fig. 2). Mitochondrial objects were segmented in Imaris to quantify object volume, surface area, sphericity, and longest-axis length (Fig. 1G-J). The interpretation of each morphological parameter is summarized in (SI Table 1). Longest-axis length describes the extent of individual segmented objects but does not measure mitochondrial connectivity or network topology. Although injury was directed at UV cone photoreceptors in the outer retina, mitochondrial changes were already detectable in inner and intermediate Müller glial compartments at 24hpt. Here, mitochondrial surface area and volume were increased in the GCL and INL, whereas the longest-axis length was increased in the GCL, IPL, INL and OPL (Fig. 1D,G,H,J). No corresponding changes in surface area or volume were detected in the IPL, OPL or ONL at this time point (Fig. 1G,H). Sphericity was increased only in the GCL and OPL, indicating that early morphological responses differed between Müller glial compartments and could not be described solely as mitochondrial elongation (Fig. 1I). By 48hpt, mitochondrial remodeling was broadly detected across the retina. Surface area and volume were increased in the GCL, IPL, OPL and ONL, while longest-axis length was increased in all five retinal layers (Fig. 1E,G,H,J). In contrast, sphericity was not significantly altered in any layer at this time point (Fig. 1I). Notably, mitochondria within the INL exhibited increased longest-axis length without corresponding changes in surface area or volume, indicating that individual morphological parameters were differentially regulated within the Müller glial cell body compartment. By 60hpt, increased longest-axis length persisted in all five retinal layers (Fig. 1J) and increased surface area persisted in the GCL, IPL, OPL and ONL (Fig. 1H), whereas increased volume was retained only in the ONL (Fig. 1G). Sphericity was increased only in the OPL (Fig. 1I).

**Figure. 1.**
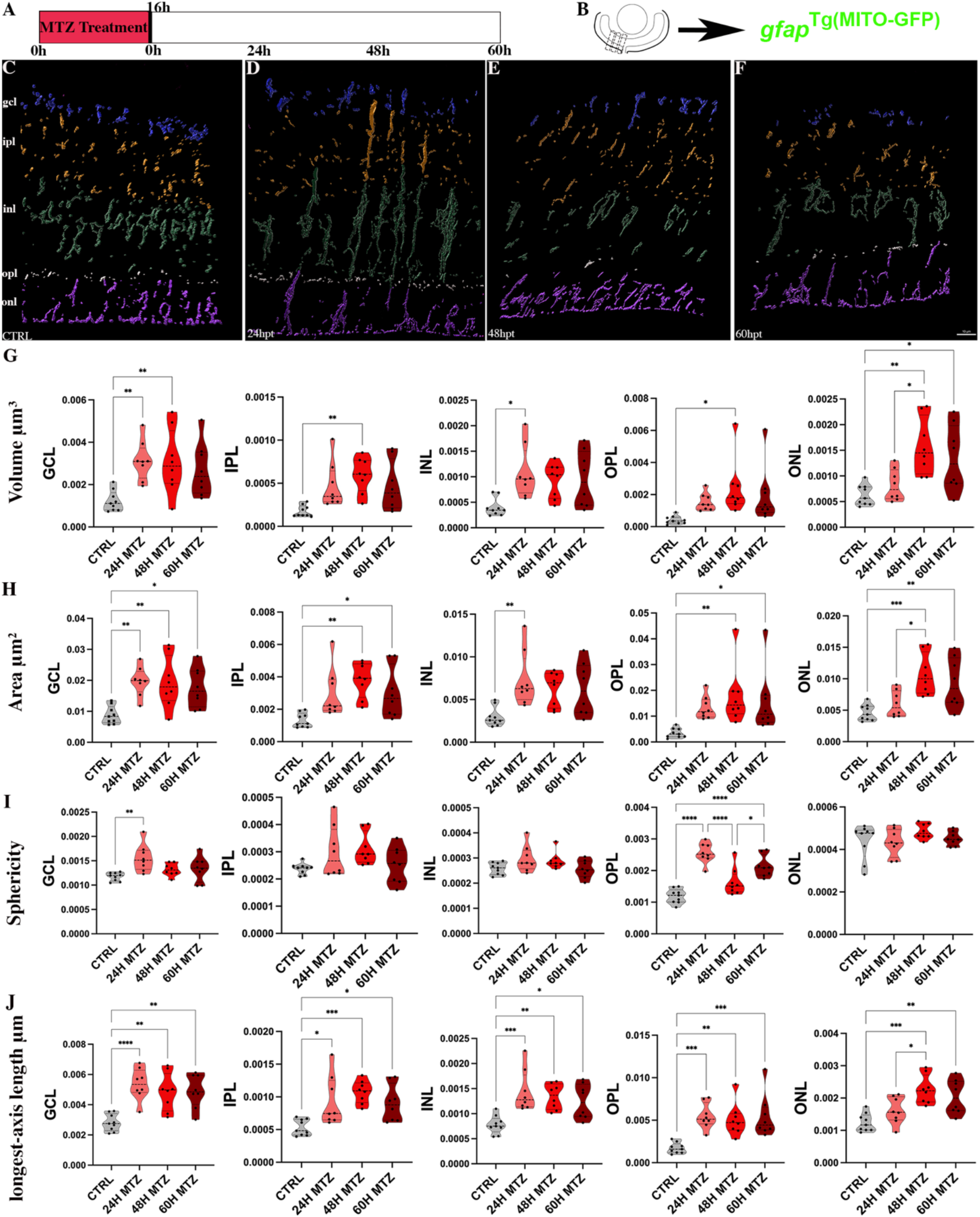
Photoreceptor injury induces spatially and temporally patterned mitochondrial remodeling in Müller glia. (A) Experimental design. Adult *gfap*^Tg(MITO-GFP)^;Tg(*sws1:nfsB-mCherry)* zebrafish were exposed to 10mM metronidazole (MTZ) for 16h to selectively ablate ultraviolet-sensitive cone photoreceptors. Müller glial mitochondrial morphology was imaged at 24, 48, and 60 h after MTZ treatment and compared with untreated controls. (B) Representative retinal image indicating the central retinal region used for mitochondrial imaging and quantitative analysis. For each biological sample (i.e. retina), three central retinal sections were imaged and from each section, three adjacent high-resolution images quantified and averaged to generate one value per fish. CTRL, *n* = 9; 24, 48 and 60hpt *n* = 8. (C-F) Representative three-dimensional reconstruction of mito-GFP-positive mitochondrial objects segmented in Imaris. Mitochondrial objects in retinal layers are pseudocolored as follows: GCL = blue; IPL = orange; INL = green; OPL = white; ONL = magenta. Segmentation and morphometric analyses were performed separately within each retinal layer. G-J) Quantification of mitochondrial-object: G) volume; H) surface area; I) sphericity and J) longest-axis length within each retinal layer. Surface area, volume, and longest-axis length are reported in µm², µm³, and µm respectively; sphericity is dimensionless. Data are presented as violin plots with the median and quartiles indicated; data points presented as individual biological samples. Statistical analyses were performed separately for each morphometric parameter and retinal layer using one-way ANOVA followed by Tukey’s multiple-comparisons test. The *P* values shown are multiplicity-adjusted. \**P* < 0.05, \*\**P* < 0.01, **\*\*\****P* < 0.001, and \*\*\*\**P* < 0.0001. Exact adjusted *P* values are provided in the SI stats table 1. Scale bar = 10 µm.

Photoreceptor injury therefore elicited a spatially and temporally patterned mitochondrial response throughout Müller glia. Changes were detectable in inner retinal compartments at 24hpt, whereas sustained increases in mitochondrial volume emerged later in processes adjacent to the injured photoreceptors. The relationships among longest-axis length, volume, and sphericity differed by compartment and time point.

### Müller glial mount a staged mitochondrial transcriptional response to injury

To define the molecular response accompanying injury-induced mitochondrial remodeling, we performed RNA sequencing of FACS-isolated Müller glia at 12, 24, 36, 48, and 60hpt (Fig. 2A; SI Fig. 3). Differentially expressed genes were identified relative to uninjured controls using an adjusted *P* value of <0.05 and an absolute fold change of >1.5. Consistent with previous studies [12, 13], our data identified distinct early, intermediate, and late injury-induced Müller glial transcriptional responses. The early response at 12hpt was dominated by reduced gene expression, shifting to gene induction during the intermediate 24–48hpt interval, and then returning to predominantly reduced expression at 60hpt (SI Fig. 4A and Supplementary Tables 2,3).

**Figure 2.**
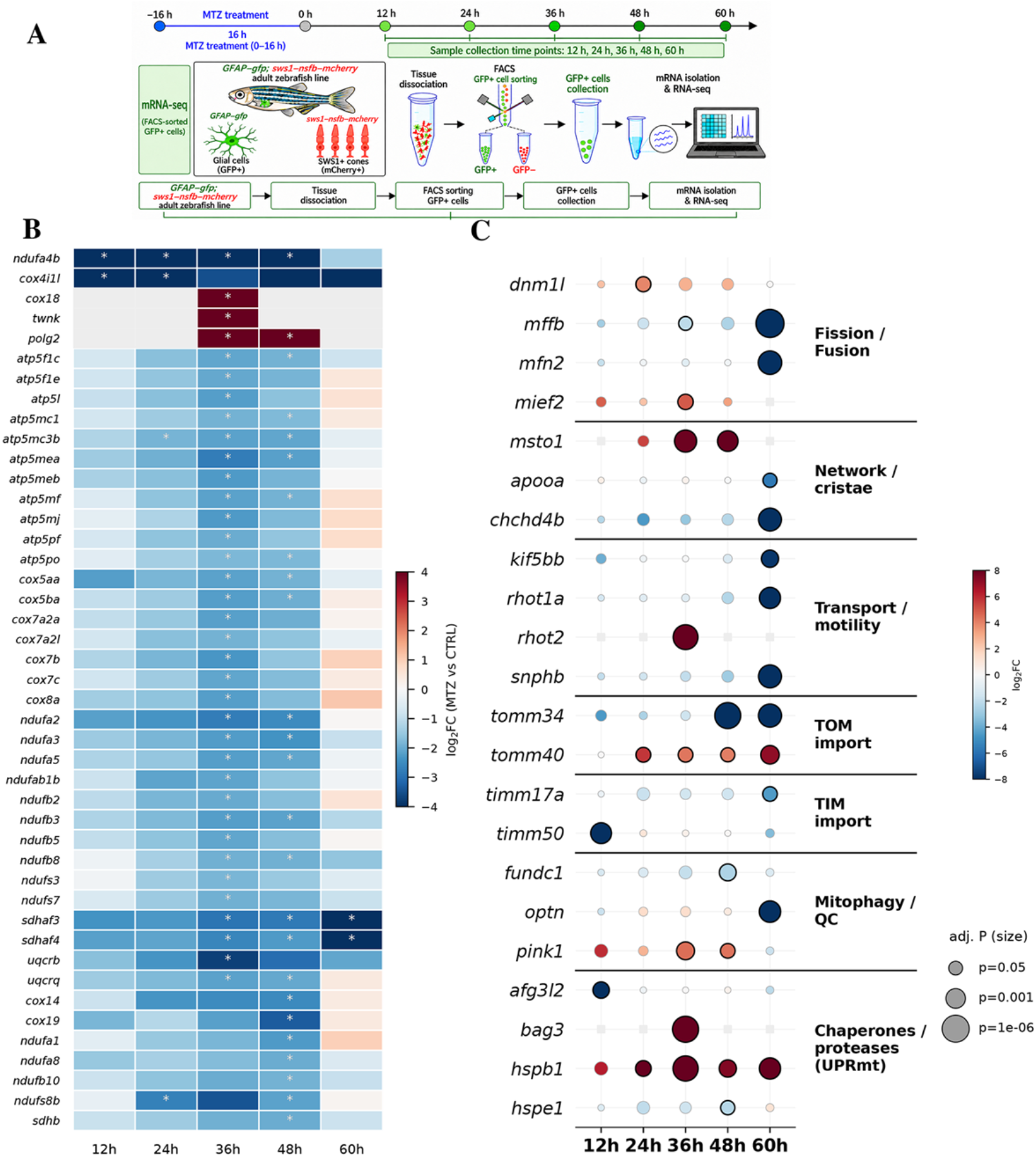
Müller glia exhibit a staged transcriptional response to photoreceptor injury. (A) Experimental design and workflow. Ultraviolet-sensitive cone photoreceptors were ablated in adult Tg(*gfap*:GFP);Tg(*sws1*:nfsB-mCherry) zebrafish. Retinae were collected at 12, 24, 36, 48, and 60hpt dissociated, and subjected to fluorescence-activated cell sorting (FACS). GFP-positive Müller glia were collected, followed by mRNA isolation and RNA sequencing. (B) Heatmap showing temporal changes in selected genes associated with mitochondrial respiratory chain function, mitochondrial assembly and maintenance. Differentially expressed genes were identified using an adjusted *P* value of < 0.05 and an absolute fold change of > 1.5. Asterisks identify genes that were significantly differentially expressed at the indicated time point (Benjamini–Hochberg-adjusted *P* < 0.05). (C) Dot plot showing the temporal expression profiles of selected genes associated with mitochondrial fission and fusion, cristae and network organization, mitochondrial transport and motility, TOM-and TIM-dependent protein import, mitophagy and mitochondrial quality control, and mitochondrial chaperone and protease responses. Dot color denotes the DESeq2 log2 fold change relative to untreated controls: red denotes increased expression and blue denotes reduced expression. Dot size reflects the adjusted *P* value, with larger dots indicating smaller *P* values. Differential-expression analyses were performed independently at each time point using DESeq2, and *P* values were adjusted for multiple testing using the Benjamini–Hochberg procedure. Complete differential-expression results and exact adjusted *P* values are provided in SI tables 2,3. in the source data.

Despite the predominance of gene induction during the intermediate injury response, respiratory-chain transcripts were broadly reduced (Fig. 2B). The respiratory chain-associated gene *ndufa4b* was reduced from 12-48hpt, while *cox4i1l* was reduced at 12 and 24hpt. By 36–48hpt, reduced expression extended to multiple nuclear-encoded components of oxidative phosphorylation complexes I–V, including members of the *nduf*, *sdh*, *uqcr*, *cox*, and *atp5* families. During the same interval however, genes associated with respiratory chain assembly and mitochondrial genome maintenance were induced. For example, *cox18* and *twnk* were induced at 36hpt, and *polg2* was induced at 36 and 48hpt. Thus, the intermediate injury-induced transcriptional response in Müller glia was characterized by broad reduction of respiratory chain subunit transcripts accompanied by selective induction of genes associated with mitochondrial assembly and maintenance. Gene-set enrichment analyses supported a reduction in respiratory-program transcripts. The KEGG oxidative phosphorylation pathway showed significant negative enrichment during the 24–48hpt interval (SI Fig. 4B). Reactome and Gene Ontology analyses likewise identified negative enrichment of gene sets associated with respiratory electron transport, ATP synthesis, and mitochondrial respiratory chain assembly (SI Fig. 4C). These pathway-level patterns occurred alongside selective induction of individual assembly and maintenance genes.

Metabolic transcripts showed mixed, time-dependent changes rather than uniform induction of glycolysis-associated genes (SI Fig. 5). For example, the glycolytic regulator *hk2* was induced at 24 and 48hpt, and *pfkfb2a* was induced at 48hpt. Conversely, *gapdhs* and *tpi1b* were reduced during the 36–48 interval, while *pfkfb4b* was reduced at several time points. Genes controlling mitochondrial pyruvate utilization were also differentially regulated: *pdk2b* was induced at 36hpt, while the mitochondrial pyruvate carrier *mpc2a* and the pyruvate dehydrogenase subunit *pdha1b* were reduced at 48hpt. Within the tricarboxylic acid and associated metabolic pathways, *aco1* was induced from 24 through 48hpt, while other genes displayed time-restricted increases or decreases.

Genes associated with mitochondrial dynamics and transport also showed distinct temporal responses (Fig. 2C). For example, the fission-associated GTPase *dnm1l* was induced at 24hpt, whereas *mief2* was induced and *mffb* was reduced at 36hpt. The mitochondrial network and cristae regulator *msto1* was induced at 36 and 48hpt. Genes associated with mitochondrial transport components were also altered, with induction of *rhot2* at 36hpt followed by reduced expression of *rhot1a*, *kif5bb*, and *snphb* at 60hpt. Additionally, the late transcriptional response included reduced expression of *mffb* and the fusion-associated gene *mfn2*. Individual components of mitochondrial protein import and quality control were differentially regulated during the same interval (Fig. 2C). *timm50* was reduced at 12hpt, whereas *tomm40* was induced from 24-60hpt. The mitochondrial quality-control kinase *pink1* was induced at 36 and 48hpt, while the mitophagy-associated receptor *fundc1* was reduced at 48hpt.

Together, the RNA sequencing data reveal a temporally staged mitochondrial response accompanying Müller glial activation. Early suppression of selected mitochondrial genes was followed during the 24–48hpt interval by coordinated reduction of oxidative phosphorylation and respiratory chain programs, together with selective induction of genes involved in respiratory complex assembly, mitochondrial genome maintenance, dynamics, transport, and stress adaptation. The late transcriptional response at 60hpt was distinguished by reduced expression of multiple genes associated with mitochondrial dynamics, transport, import, and metabolism. These transcriptional changes place the compartment-specific mitochondrial remodeling within a broader molecular response to photoreceptor injury. We next examined the tissue-level metabolic changes accompanying this response.

### Photoreceptor injury alters the eyecup metabolic environment during Müller glia activation

Photoreceptors exchange metabolites with neighboring retinal cells, and their degeneration is associated with changes in the retinal metabolome [14, 15]. We therefore asked what tissue-level metabolic changes accompany Müller glial activation after selective photoreceptor loss. We performed untargeted metabolomic profiling of whole eyecups from uninjured controls and at 12, 24, 36, 48, and 60hpt (SI Fig. 6; Supplementary Table 4). These measurements capture changes across the retina and associated tissues, providing the tissue metabolic context for the transcriptional response in isolated Müller glia.

Central-carbon metabolites changed over the injury time course (Fig. 3). Glucose 6-phosphate and several carbon-and pentose-associated metabolites were reduced during the early response, whereas pyruvate increased at 36hpt onward and α-ketoglutarate also increased during the intermediate response. These changes coincided with altered expression of central-carbon and mitochondrial genes in sorted Müller glia (Figs. 2,4). The concurrence of these changes highlights pyruvate handling during the period of mitochondrial remodeling, although metabolite abundance and transcript levels do not establish the direction of metabolic flux.

**Figure 3.**
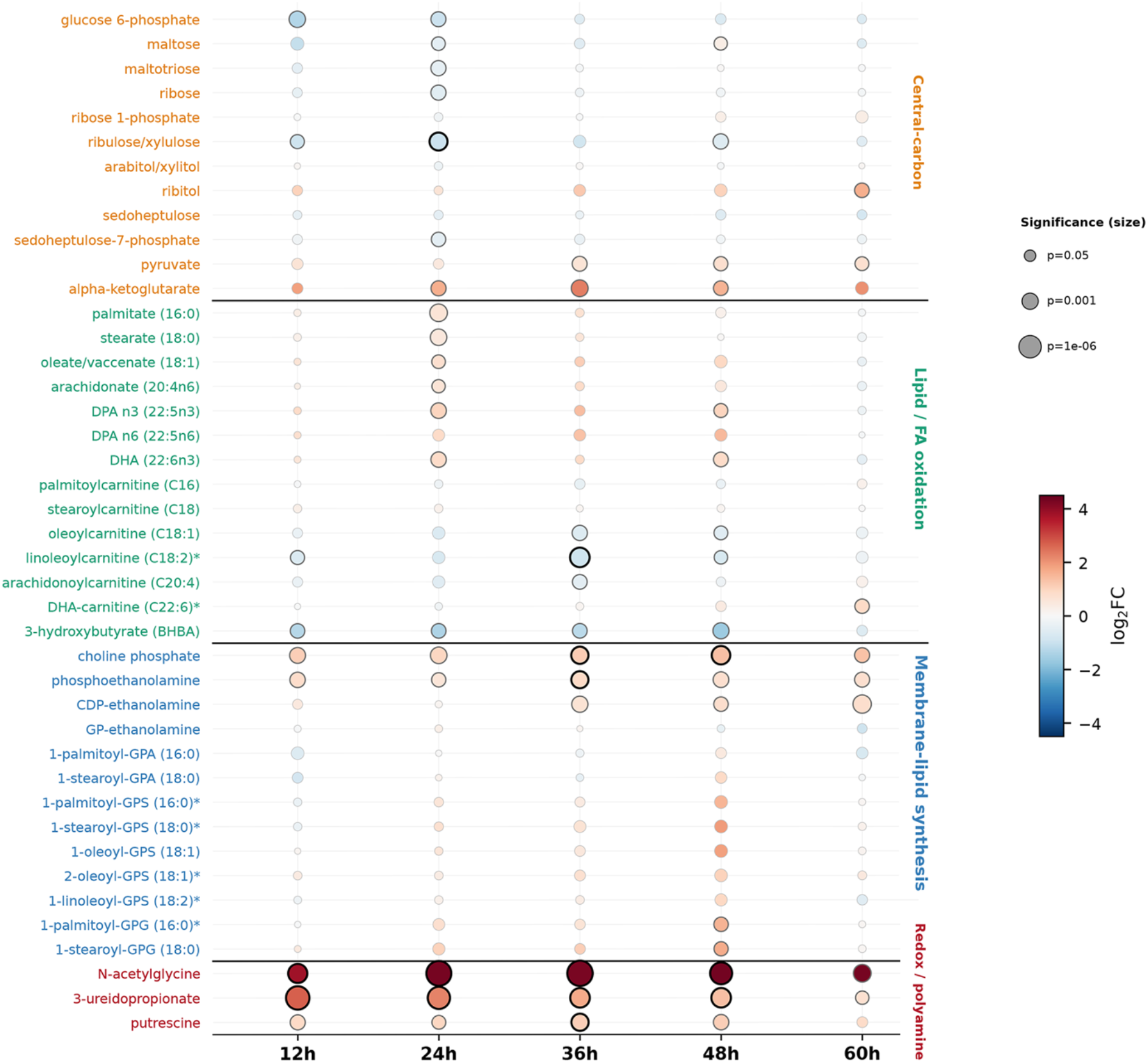
Photoreceptor injury induces temporally patterned metabolic changes in the eyecup. Untargeted metabolomic profiling of whole eyecups collected at 0 (uninjured control), 12, 24, 36, 48, and 60hpt (n=4 biological replicates per time point). The dot plot shows selected metabolites grouped by their associations with central carbon metabolism, lipid metabolism and fatty acid oxidation, membrane-lipid synthesis, and redox and polyamine metabolism. Dot color indicates the log2 fold change in metabolite abundance relative to untreated controls: red denotes increased abundance and blue denotes reduced abundance. Dot size reflects the nominal *P* value, with larger dots indicating smaller *P* values. Black outlines denote changes significant after false discovery rate correction (*q* < 0.05). MTZ-treated and untreated samples were compared separately at each time point using Welch’s two-sample *t* test on natural-log-transformed scaled intensity values. Asterisks identify metabolites assigned with high confidence by Metabolon but not confirmed using authentic chemical standards. Exact *P* and *q* values and the complete metabolomics dataset are provided in SI table 4.

Lipid-associated metabolites were similarly remodeled after retinal injury (Fig. 3). Multiple long-chain free fatty acids, including palmitate, stearate, oleate/vaccenate, arachidonate, docosapentaenoate, and docosahexaenoate increased at 24hpt, followed by reductions in several long-chain acylcarnitines at 36–48hpt. Metabolites associated with membrane-lipid synthesis also changed (Fig. 3). Choline phosphate and phosphoethanolamine increased across the time course, while CDP-ethanolamine and selected lysophospholipids increased at later time points. Changes in membrane-lipid-associated metabolites occurred during the interval of mitochondrial remodeling (Fig. 1) and coincided with altered expression of lipid handling genes in Müller glia (Fig. 4).

**Figure 4.**
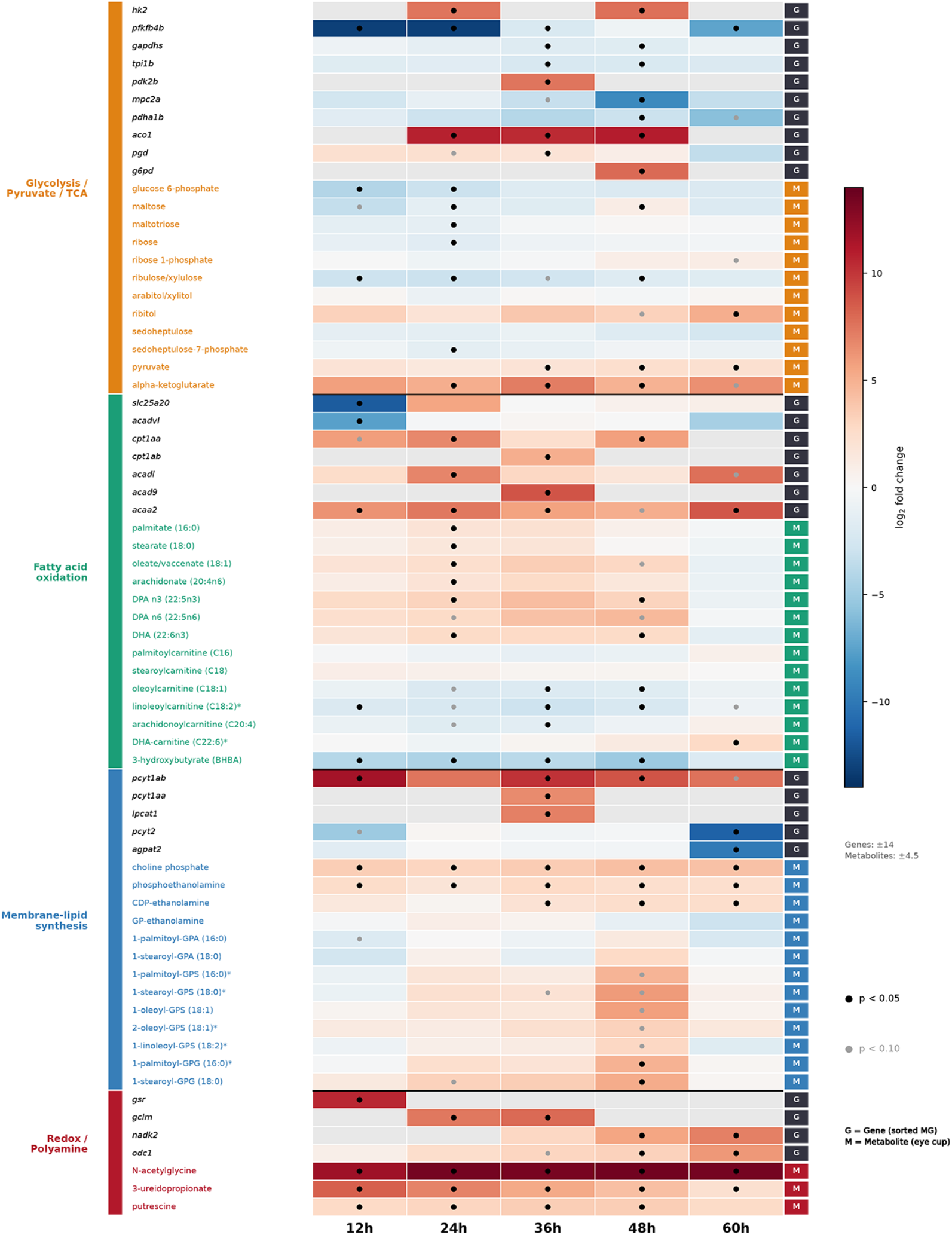
Temporal alignment of Müller glial gene expression changes and eyecup metabolite changes after photoreceptor loss. Heatmap aligning changes in selected genes and metabolites associated with glycolysis/pyruvate metabolism and the TCA cycle, fatty acid oxidation, membrane-lipid synthesis, and redox/polyamine metabolism at 12, 24, 36, 48, and 60hpt. Gene expression measurements were obtained from FACS-isolated Müller glia, and metabolite measurements from whole eyecups. Values represent log₂ fold changes relative to time-matched uninjured controls; red denotes increased expression or abundance and blue denotes reduced expression or abundance. Gene and metabolite values were displayed using separate ranges (genes: −14 to +14; metabolites: −4.5 to +4.5) to permit visualization within the same matrix. Black circles indicate *P* < 0.05, and gray circles indicate 0.05 ≤ *P* < 0.10, using adjusted P values for genes and nominal P values for metabolites. Asterisks indicate metabolites annotated without confirmation using authentic chemical standards. The aligned datasets illustrate temporal correspondence but do not establish Müller glia-specific metabolite changes or metabolic flux.

The metabolic response extended beyond central carbon and lipid metabolism (Fig. 3). Additional eyecup changes included a sustained increase in N-acetylglycine and a transient rise in putrescine. The rise in putrescine preceded induction of *odc1* and the later appearance of DNA replication and cell cycle-associated transcripts in Müller glia (Fig. 4). The temporal relationship between putrescine accumulation and cell cycle gene induction is consistent with remodeling of polyamine metabolism during Müller glial activation, although the cellular origin of putrescine cannot be determined from our whole eye measurements.

Together, these data identify temporally patterned changes in central carbon, lipid, and polyamine-associated metabolites after photoreceptor loss. These tissue-level changes overlap with mitochondrial remodeling and metabolic gene regulation in Müller glia, although individual metabolite changes cannot be assigned to Müller glia cells (Fig. 4). Pyruvate accumulation, coupled to the altered expression of genes regulating mitochondrial pyruvate utilization, prompted us to test whether perturbing the PDK–PDH axis alters the Müller glial injury response.

### DCA treatment alters PDHA1 phosphorylation and outer-retina mitochondrial remodeling during the injury response

To test whether perturbing the PDK–PDH axis alters the Müller glial injury response, we ablated UV cones and treated with dichloroacetate (DCA), a well-known inhibitor of PDK activity [16]. PDKs phosphorylate PDHA1, inhibiting the PDH complex; we therefore examined phospho-PDH1A as a biochemical readout of PDK–PDH axis regulation in the injured retina. DCA treatment markedly reduced phospho-PDHA1 immunoreactivity relative to MTZ treatment alone, consistent with engagement of this regulatory axis in vivo (SI Fig. 7). The extent of MTZ-induced UV cone loss did not differ detectably between MTZ and MTZ+DCA groups (SI Fig. 8).

DCA treatment alone produced no overt mitochondria morphology changes in the absence of UV cone ablation (SI Fig. 9); however, in conjunction with UV cone ablation, DCA altered mitochondrial morphology in a compartment- and time-dependent manner (Fig. 5, SI Fig. 10). By 48hpt, DCA treatment reduced mitochondrial surface area, volume, and longest-axis length in the ONL, indicating attenuation of the injury-associated expansion and elongation of mitochondrial objects within Müller glial processes adjacent to the damaged photoreceptors (Fig. 5D,G,I). DCA also reduced sphericity in the INL at 24 and 48hpt and increased sphericity in the OPL at 48hpt (Fig. 5C,D,F,G,I). At 60hpt, reduced mitochondrial surface area and volume persisted in the ONL, whereas effects on longest-axis length and sphericity were no longer detected (Fig. 5E,H,I). Other Müller glial compartments showed no corresponding changes in mitochondrial surface area, volume, or longest-axis length (SI Fig. 11).

**Figure 5.**
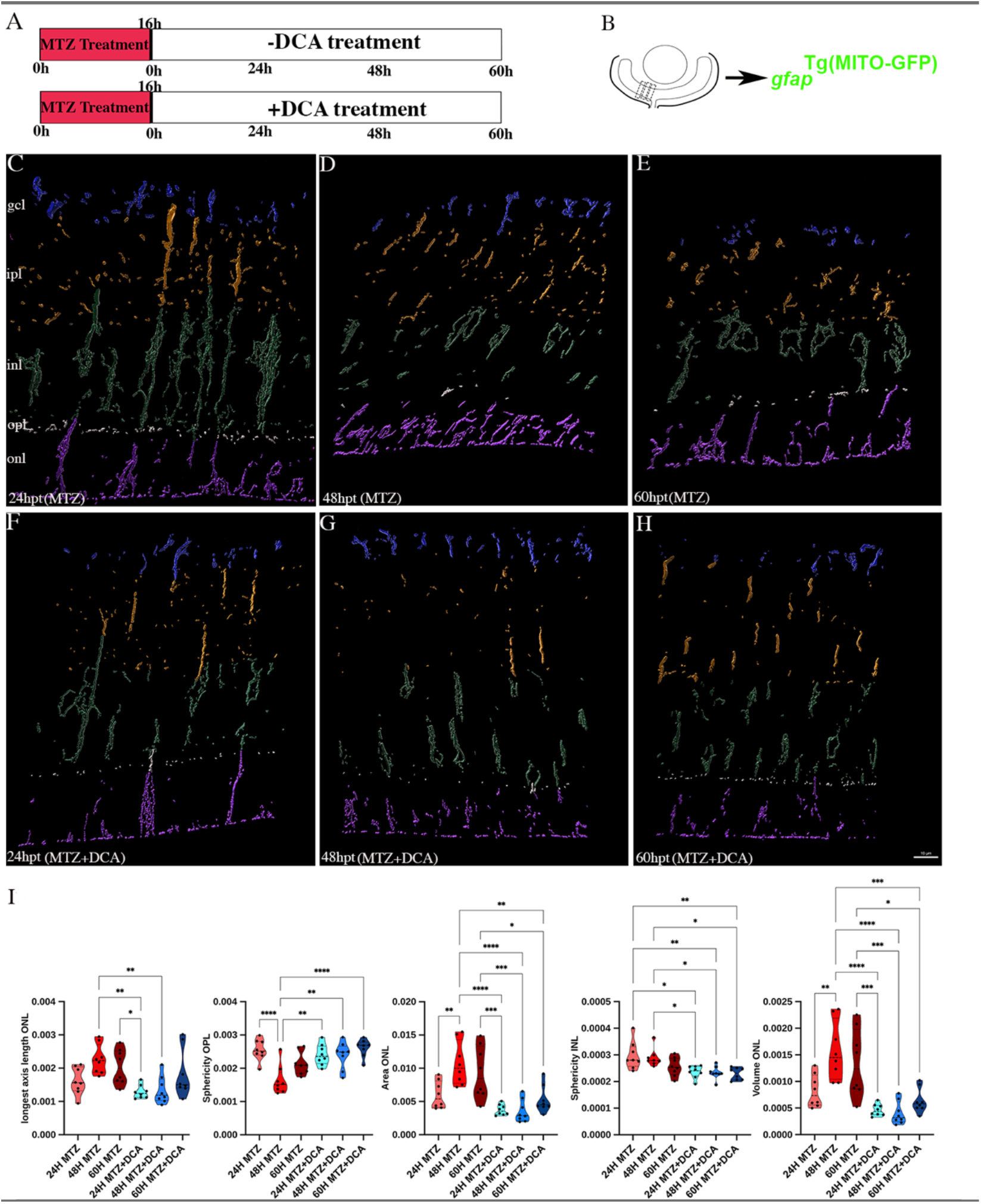
DCA attenuates injury-associated mitochondrial remodeling in outer-retinal Müller glial processes. (A) Experimental design. (B) Representative retinal image indicating the central retinal region used for mitochondrial imaging and quantitative analysis. For each biological sample (i.e. retina), three central retinal sections were imaged and from each section, three adjacent high-resolution images quantified and averaged to generate one value per fish. *n* = 8 for each condition. (C–H) Representative three-dimensional reconstruction of mito-GFP-positive mitochondrial objects segmented in Imaris. Mitochondrial objects in retinal layers are pseudocolored as follows: GCL = blue; IPL = orange; INL = green; OPL = white; ONL = magenta. Segmentation and morphometric analyses were performed separately within each retinal layer. (I) Quantification of mitochondrial-object surface area, volume, longest-axis length and sphericity within each layer. Surface area, volume, and longest-axis length are reported in µm², µm³, and µm respectively; sphericity is dimensionless. Graphs show the mitochondrial-object parameters for which significant time-matched differences between MTZ and MTZ+DCA treatments were detected: surface area and volume in the ONL, sphericity in the OPL and INL, and bounding-box longest-axis length in the ONL. Data are presented as violin plots with the median and quartiles indicated; data points presented as individual biological samples. Statistical analyses were performed separately for each morphometric parameter and retinal layer using one-way ANOVA followed by Tukey’s multiple-comparisons test. *P* values shown are multiplicity-adjusted. \**P* < 0.05, \*\**P* < 0.01, **\*\*\****P* < 0.001, and \*\*\*\**P* < 0.0001 Exact adjusted *P* values and all additional comparisons that resulted in non-significant differences are provided in SI stats table 2. Scale bar =10 µm.

These findings demonstrate that injury-associated mitochondrial remodeling is sensitive to perturbation of the PDK–PDH axis, with sustained DCA-associated reductions in mitochondrial size detected specifically in ONL Müller glial processes adjacent to injured photoreceptors. Together, these results identify a spatially patterned mitochondrial response to metabolic perturbation during Müller glial activation.

### DCA treatment attenuates Müller glial reprogramming and proliferation

Having established that DCA altered retinal PDHA1 phosphorylation and Müller glial mitochondrial remodeling, we next asked whether treatment also affected progression toward reprogramming and proliferative activation. We compared transcriptomes of FACS-isolated Müller glia from MTZ- and MTZ+DCA-treated retinas at 12, 24, 36, 48, and 60hpt. The transcriptional effects of DCA changed over the injury-response time course (SI Fig. 12A; Supplementary Table 5). Early effects included increased expression of chromatin associated genes at 12hpt and reduced expression of ribosome associated genes at 24hpt. By 48hpt, DCA-responsive genes converged on processes required for proliferative entry. Gene set enrichment analysis identified coordinated reductions in programs associated with cell-cycle regulation, mitotic progression, chromosome segregation, and DNA metabolism. Suppression of this program persisted at 60hpt, when DNA replication and DNA templated replication pathways were reduced (SI Fig. 12B). Thus, PDK inhibition altered the Müller glial transcriptional trajectory before the peak proliferative response.

These reductions occurred alongside selective changes in metabolic and mitochondrial gene expression. At 48hpt, DCA increased expression of multiple glycolysis-associated genes, including *ldha*, *gpia*, *pgk1*, *eno1a*, *pgam1a*, and *pfkfb3*, while reducing transcripts associated with mitochondrial carbon metabolism and respiratory chain function, including *ogdhl*, *me3*, *cox10*, and *etfb* (SI Fig. 12C). By 60hpt, expression of multiple respiratory chain and ATP-synthase components was reduced, whereas genes associated with lipid biosynthesis, mitochondrial biogenesis and dynamics, and autophagy were increased. These included *srebf2*, *cers5*, *sgms1a*, *elovl6*, *ppargc1a*, *nrf1*, *mfn2*, *mffb*, *ulk2*, and *map1lc3a*. Notably, *pdk2a* was strongly induced at 60hpt despite pharmacological PDK inhibition, consistent with a potential compensatory response within the PDK–PDH regulatory axis (SI Fig. 12D). DCA therefore suppressed proliferation-associated programs while producing time dependent increases and decreases in metabolic and mitochondrial transcripts, reshaping rather than uniformly dampening the Müller glial injury response.

The transcriptional effects extended to individual regulators and markers of Müller glial regeneration. Expression of the regeneration-associated marker *tuba1a* [17] and the DNA replication factor *mcm4* was reduced by 48hpt (Fig. 6A), coincident with pathway-level suppression of cell cycle and DNA metabolism programs. By 60hpt, DCA also reduced expression of *ascl1a*, a central regulator of Müller glial reprogramming [18], together with *pcna* and *cdk1* (Fig. 6A).

**Figure 6.**
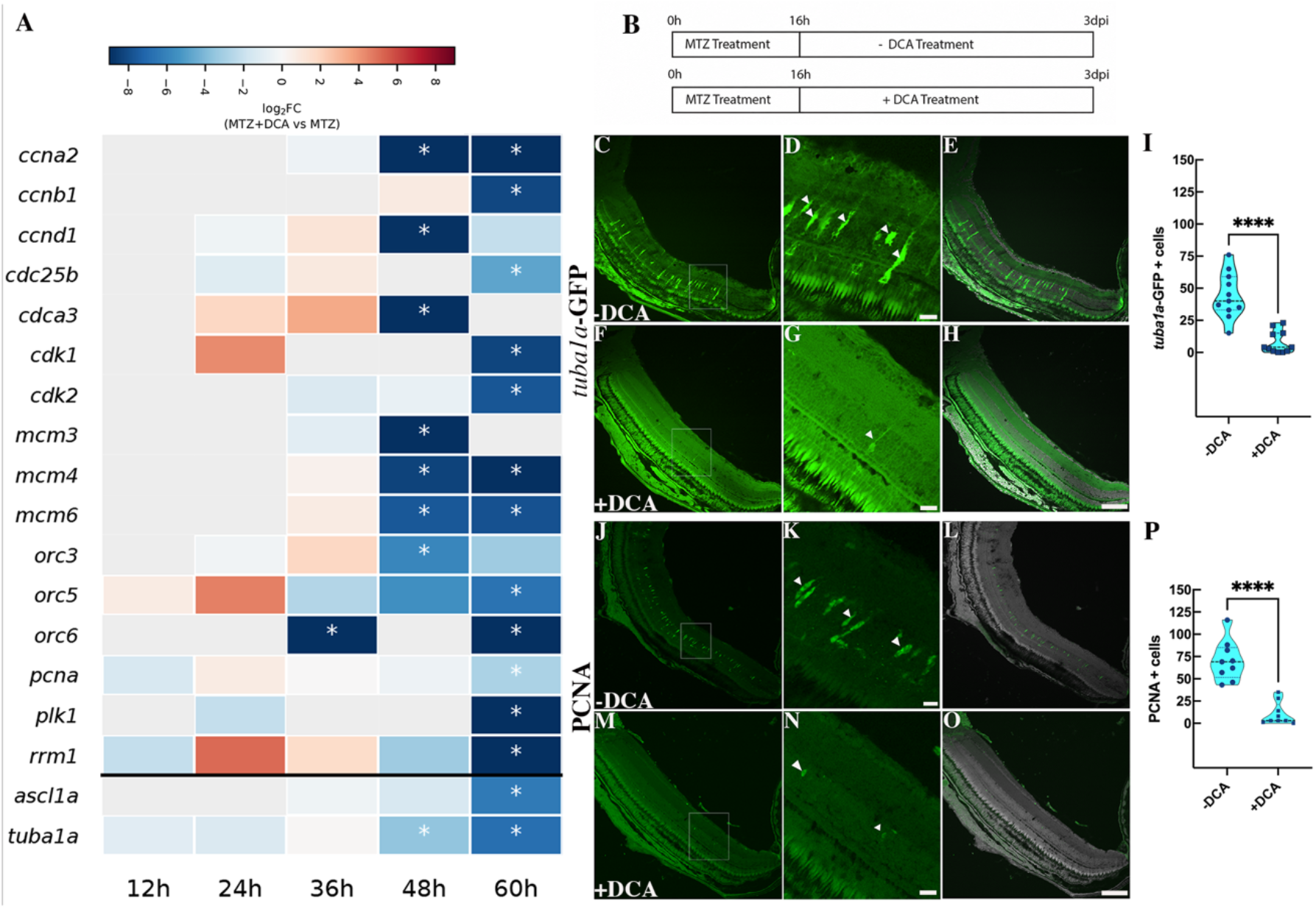
DCA suppresses Müller glia reprogramming-associated gene expression and proliferative activation. (A) Heatmap showing changes in expression of selected regeneration- and cell-cycle-associated genes in FACS-isolated Müller glia at 12, 24, 36, 48, and 60hpt. Colors indicate the DESeq2 log2 fold changes in MTZ+DCA-treated samples relative MTZ-treated) samples; red denotes increased expression and blue denotes reduced expression. Asterisks denote significant differential expression at the indicated time point (Benjamini– Hochberg-adjusted *P* < 0.05). (B) Experimental design. (C–H) Representative retinal sections from MTZ-treated (C–E) and MTZ+DCA-treated (F–H) Tg(*tuba1a:GFP)* fish at 3dpi. C and F show low-magnification views; D and G show enlargements of the boxed regions, and E and H show merged DAPI images. Arrowheads indicate *tuba1a*:GFP-positive Müller glia. (I) Quantification of *tuba1a*:GFP-positive cells in MTZ-and MTZ+DCA-treated retinae. (J–O) Representative PCNA-immunolabeled retinal sections from MTZ-treated (J–L) and MTZ+DCA-treated (M–O) fish at 3dpi. J and M show low-magnification views; K and N show enlargements of the boxed regions, and L and O show the corresponding PCNA and DAPI merged images. Arrowheads indicate PCNA-positive cells. (P) Quantification of PCNA-positive cells per retina. Each point in I and P represents one biological sample (retina); n=11 for Tg(*tuba1a:GFP)* and n=9 for PCNA. Violin plots show the median and quartiles. Statistical significance was evaluated using two-tailed Mann–Whitney U test. \*\*\*\**P* < 0.0001. Exact *P* values are provided in SI stats table 3. Scale bar = 50 µm.

Finally, we addressed whether these transcriptional changes were accompanied by an impairment of Müller glial reprogramming and proliferative activation (Fig. 6B). At 3 days post injury (dpi), photoreceptor ablation induced robust *tuba1a*:GFP expression in Müller glia (Fig. 6E-G) and accumulation of PCNA^+^ cells (Fig. 6J-L). DCA treatment reduced *tuba1a*:GFP^+^ Müller glia by approximately 90% (Fig. 6F-I; P < 0.0001) and PCNA^+^ cells by approximately 95% (Fig. 6M–P; *P* < 0.0001). Together, these results demonstrate that the DCA-dependent transcriptional suppression of reprogramming and cell cycle genes was accompanied by marked suppression of reprogramming-associated reporter expression and proliferative activation after photoreceptor injury.

## Discussion

Here, we demonstrate that localized photoreceptor loss elicited a distributed, compartment-specific mitochondrial response in zebrafish Müller glia. Changes were detectable in inner retinal compartments before sustained outer retinal remodeling, showing that the response was not confined to processes adjacent to damaged photoreceptors. These structural changes accompanied selective mitochondrial gene regulation and tissue-level metabolic changes during Müller glial activation. DCA reduced retinal PDHA1 phosphorylation, attenuated outer retinal mitochondrial remodeling, and markedly suppressed reprogramming-associated and proliferative responses. Together, these findings establish mitochondrial remodeling within the early Müller glial regenerative response and identify the PDK–PDH axis as a critical component of regenerative activation.

Müller glia are highly polarized cells whose endfeet, somata and processes occupy retinal compartments with distinct requirements for local energy production, ion buffering and redox regulation, amongst others [19]. The mitochondria must therefore be organized to meet the biological demands of Müller glia during homeostatic conditions as well as during the injury and regenerative responses. Consistent with this organization, photoreceptor injury elicited different combinations of changes in mitochondrial object volume, surface area, longest-axis length, and sphericity across retinal layers. The most sustained volume increase was detected in Müller glial processes facing the damaged outer retina suggesting that mitochondrial adaptation is locally tuned to the site of photoreceptor loss. The differing combinations of morphological changes across compartments show that the response is not a uniform change in mitochondrial size or shape. However, because these measurements describe segmented objects they cannot directly establish mechanisms like increased fusion or redistribution between compartments after injury. The observed phenotype may thus reflect varying contributions from organelle growth, redistribution, turnover, and post-transcriptional control of the fission–fusion machinery that could each contribute to the observed morphologies. Determining their relative contributions will require additional high-resolution imaging analyses and cell- and potentially even layer-specific perturbations. Nonetheless, this study establishes compartment-specific mitochondrial remodeling as a novel component of the Müller glia response to injury.

The timing of mitochondrial remodeling coincided with the transition from injury response to reprogramming in zebrafish Müller glia. Zebrafish retinal regeneration proceeds through injury responsive signaling, ascl1a-dependent reprogramming of Müller glia, and production of proliferative Müller glia-derived progenitors [3, 18]. Here, morphological remodeling of mitochondria was detectable by 24hpt, became widespread by 48hpt, before the prominent proliferative response at 3dpi. The transcriptional response was similarly staged. Early reductions in selected mitochondrial transcripts were followed by broad reduction of respiratory chain genes alongside the selective induction of genes associated with mitochondrial genome maintenance, respiratory complex assembly, protein import and mitochondrial quality control. Rather than uniformly increasing or decreasing in expression, these data suggest that Müller glia reorganize distinct aspects of mitochondrial function as they respond to injury. These changes overlapped with *ascl1a* induction and were followed by increased expression of cell cycle transcripts like *cdk2, ccna2*, and *plk1*. This sequence of events places mitochondrial adaptation within the progression toward Müller glial reprogramming and cell cycle re-entry.

Indeed, the timing of mitochondrial injury responses may also be relevant to regenerative outcome. Following spinal cord injury in *Xenopus laevis*, regenerative and nonregenerative developmental stages show different mitochondrial responses; the nonregenerative stage exhibits a delayed response followed by mitochondrial dysfunction [20]. This comparison raises the possibility that when and how mitochondria remodel could matter as much as the presence of a morphological response. Anusha et al. also recently reported delayed caudal fin regeneration and abnormal mitochondrial ultrastructure in aged and rotenone-treated zebrafish, further linking mitochondrial dysfunction with impaired regeneration [21].

Interestingly, our findings align with regenerating retinal ganglion cells (RGCs) in adult zebrafish. Following optic nerve crush, there is a transient reduction in RGC dendritic mitochondria during dendritic retraction, enlarged mitochondria within the regenerating optic nerve and tract, and increased indices of mitochondrial fission and biogenesis in RGC somata; dendritic mitochondrial density later returns toward baseline as dendrites regrow [8]. Our results identify a related principle of compartment-specific mitochondrial remodeling in an injury-responsive retinal glial cell, although the compartments and regenerative programs differ between these cell types. Studies in neurons also highlight the importance of mitochondrial localization. For example, increasing mitochondrial transport through ARMCX1 promotes axonal regeneration after optic nerve crush, demonstrating that the availability of mitochondria at distant cellular compartments can influence repair independently of total mitochondrial abundance [22]. Additionally, retrograde mitochondrial transport from distal axonal compartments to the cell body was shown to be required to sustain mitochondrial DNA replication and nuclear expression of mitochondrial biogenesis genes in zebrafish neurons [23]. These studies provide a functional context for asking how mitochondrial positioning contributes to the compartment-specific Müller glial regenerative response in the zebrafish retina.

Evidence from other regenerative systems supports a functional relationship between mitochondrial state and regenerative cell identity. During planarian regeneration, depletion of the mitochondrial fusion regulator opa1 impairs neoblast pluripotency and impairs regeneration, whereas concurrent manipulation of drp1 partially restores mitochondrial organization and regenerative capacity [24]. In mouse neural stem cells, deletion of OPA1 or MFN1/2 impairs self-renewal and alters lineage decisions through ROS–NRF2-dependent signaling to the nucleus [4]. These studies demonstrate that mitochondrial dynamics can influence stem or progenitor behaviors during regenerative responses. More broadly, mitochondrial dynamics, quality control, redox signaling, and metabolite production are increasingly recognized as active regulators of tissue regeneration rather than merely downstream responses to increased energy demand [25]. Our findings extend this principle to an injury-responsive glial cell in an intact sensory tissue and add an anatomical dimension by showing that the response varies across the radial compartments of a single Müller cell. A functional connection between mitochondrial maintenance and regenerative competence has been demonstrated in zebrafish fin regeneration. Here, injury activated the Sirt1-dependent mitochondrial unfolded-protein response, and genetic or pharmacological disruption of this pathway impaired regenerative growth [26]. Sirt1 activity was linked to NAD⁺ metabolism and mitochondrial biogenesis, connecting organelle proteostasis with the metabolic state of regenerating tissue. The induction of mitochondrial maintenance and quality control genes in Müller glia may similarly represent an adaptive program that supports progression toward a regenerative state.

The mitochondrial response in Müller glia occurred as the injured eye cup underwent extensive metabolic changes. Early reductions in glucose 6-phosphate and related metabolites were followed by increased pyruvate and α-ketoglutarate, while increases in several long-chain free fatty acids preceded reductions in selected acylcarnitines. Metabolites associated with membrane-lipid synthesis also changed during this interval. In parallel, sorted Müller glia altered expression of genes associated with glycolysis, pyruvate handling, lipid metabolism, and mitochondrial maintenance. Together, these observations place Müller glial mitochondrial remodeling within a changing metabolic environment and identify several processes that may contribute to the regenerative response.

Recent work demonstrates that metabolic changes can influence cell state transitions. In the developing mouse retina, increasing glycolytic flux accelerated retinal progenitor-cell differentiation and rod photoreceptor maturation, while inhibiting glycolysis suppressed these activities, with evidence implicating Wnt signaling as a downstream mediator [27]. Glycolysis also supports FGF2-induced reprogramming of embryonic chicken retinal pigment epithelium (RPE) into neural retina [16]. Although these studies concern different ocular cell types and events, they establish that metabolic changes can influence retinal cell state transitions rather than simply accommodate their energetic requirements and provide useful context for asking whether metabolism participates in injury-dependent Müller glia reprogramming. Our metabolomic measurements capture the whole eye cup, including the neural retina, RPE, and other cells that may exchange metabolites during injury. The metabolomic and sorted-cell transcriptional datasets therefore provide complementary views of the tissue environment and Müller glial response; individual metabolite changes cannot yet be assigned to Müller glia.

The DCA experiments provide functional evidence that the regenerative response is sensitive to perturbation of the PDK–PDH regulatory axis. DCA attenuated the injury-associated increase in mitochondrial size and longest-axis length in Müller glial process in the ONL, with distinct or more limited effects in inner retinal compartments. The strong ONL response is notable because these Müller processes are adjacent to the injured photoreceptors and showed sustained mitochondrial remodeling after injury. Thus, perturbing the PDK–PDH axis changed the spatial pattern of mitochondrial remodeling without producing the same morphological response throughout Müller glia.

DCA effects extended to Müller glial activation; DCA produced time-dependent changes in metabolic and mitochondrial transcripts while suppressing reprogramming-and cell cycle-associated programs and markedly reducing *tuba1a*:GFP and PCNA readouts at 3dpi. A related result was reported during FGF2-induced reprogramming of embryonic chicken RPE: PDK inhibition reduced inhibitory PDH phosphorylation and prevented acquisition of neural retinal fate, redirecting the transcriptional response toward an epithelial to mesenchymal program [16]. Together, the two studies identify a critical role for the PDK–PDH axis during distinct retinal cell state transitions. In our experiments, the coordinated effects of DCA on phospho-PDHA1, Müller glial mitochondrial morphology, transcription, and regenerative readouts support a connection between metabolic regulation and Müller glial activation. Future studies are required to determine if the DCA-induced mitochondrial changes directly contribute to the reduced regenerative response or whether they occur alongside other consequences of DCA treatment.

Our findings place mitochondrial organization alongside transcriptional and metabolic remodeling as a component of Müller glial regenerative activation. Perturbation of the PDK–PDH axis alters mitochondrial remodeling and markedly suppresses reprogramming-associated and proliferative responses, identifying the PDK–PDH axis as a point of sensitivity during Müller glial regenerative activation. Determining which mitochondrial adaptations support this transition, and whether comparable responses are constrained in mammalian Müller glia, may help define the metabolic requirements for endogenous retinal repair.

## Materials and Methods

Detailed materials and methods are provided in **SI Appendix – Methods**.

### Zebrafish Husbandry and Ethics

Adult zebrafish (*Danio rerio*) were maintained at the University of Texas at Austin Zebrafish Facility under standard husbandry conditions. Fish were housed in a recirculating aquatic system at 28°C under a 12-hour light/12-hour dark photoperiod. The wild-type AB strain was used for all experiments. Transgenic lines, including *gfap*^Tg(MITO-GFP)^, Tg(*tuba1a*:GFP), and *Tg*(*gfap*:GFP;*sws1*:nsfb-mCherry), were maintained on the AB genetic background. All experimental procedures were approved by the Institutional Animal Care and Use Committee (IACUC) at the University of Texas at Austin and were conducted in accordance with institutional and federal guidelines for the care and use of laboratory animals.

### Generation of *gfap*^Tg(MITO-GFP)^

*gfap*^Tg(MITO-GFP)^ was generated using a CRISPR/Cas9-mediated intron-targeting strategy as previously described [28]. This approach targets the final intron of the endogenous *gfap* locus while preserving the endogenous coding sequence and regulatory elements. The sgRNA sequence used for genome editing was 5′-GTGCGCAACACATAGCACCA-3′. The donor plasmid was constructed using the backbone from Addgene plasmid #65564.

### Photoreceptor Ablation and Dichloroacetate (DCA) Treatment

Selective UV cone photoreceptor ablation was induced using the nitroreductase/metronidazole (NTR/MTZ) system in adult Tg(*gfap*:GFP;*sws1*:nsfb-mCherry), *gfap*^Tg(MITO-GFP)^;*sws1*:nsfb-mCherry and Tg(*tuba1a*:GFP;*sws1*:nsfb-mCherry) zebrafish (3–5 months of age). Photoreceptor ablation was induced by immersing fish in system water containing 10mM metronidazole (MTZ; Sigma-Aldrich), freshly prepared immediately before each experiment. Fish were exposed to MTZ for 16h at 28°C. For metabolic intervention experiments, sodium dichloroacetate (DCA; Sigma-Aldrich, 347795) was dissolved directly in system water and administered at a final concentration of 70mM post MTZ treatment.

### Bulk RNA Sequencing of FACS-Sorted Müller Glia

To quantify the transcriptional response of Müller glia following photoreceptor ablation, GFP-positive Müller glia were isolated Tg(*gfap*:GFP;*sws1*:nsfb-mCherry), by fluorescence-activated cell sorting (FACS) at six experimental time points: uninjured control and 12, 24, 36, 48, and 60 hours post-injury (hpt). For each biological replicate (n = 3 per time point), six retinae were collected from three adult zebrafish (3–5 months of age), pooled and enzymatically dissociated.

Retinal cells isolated from age-matched wild-type (non-transgenic) zebrafish were used to establish background fluorescence and define GFP-positive sorting gates. GFP-positive Müller glia were sorted directly into SMART-Seq HT lysis buffer (Takara Bio). Full-length cDNA was synthesized using the SMART-Seq HT kit (Takara Bio) Sequencing libraries were prepared at the UTHealth Houston Genomics Core using the Illumina Nextera XT DNA Library Preparation Kit (Illumina). Paired-end sequencing (2 × 100 bp; 200-cycle chemistry) targeting approximately 20 million reads per sample.

### Immunohistochemistry

Adult zebrafish (3–5 months of age) were euthanized, and eyes were dissected for cryosectioning. For proliferating cell nuclear antigen (PCNA) and *Tg*(*tuba1a*:GFP) experiments, eyes were dissected in phosphate-buffered saline (PBS) and fixed overnight in 4% paraformaldehyde (PFA) at 4°C. For phospho-pyruvate dehydrogenase E1α (Ser293) [phospho-PDHα1 (Ser293)] immunostaining, all procedures were performed in the presence of phosphatase inhibitors to preserve protein phosphorylation.

### Sample Preparation and Immunolabeling for NSPARC Super-Resolution Imaging

To visualize mitochondrial morphology in GFAP-expressing cells at super-resolution, retinae were dissected from adult *gfap*^Tg(MITO-GFP)^ zebrafish (3–5 months of age) in ice-cold oxygenated Ames’ medium and processed for immunolabeling followed by epoxy resin embedding. Samples were labeled using a conventional primary/secondary antibody strategy for Nikon Spatial Array Confocal (NSPARC) imaging.

### Whole eye cup Metabolomics

To characterize metabolic changes associated with retinal regeneration, global untargeted metabolomic profiling was performed on eye cup collected from adult zebrafish (3–5 months of age) at six experimental time points: uninjured control and 12, 24, 36, 48, and 60 hours post-injury (n = 4 biological replicates per time point). Approximately 30 mg of retinal tissue was collected for each biological replicate.

### Image acquisition, quantification and statistics

*gfap*^Tg(MITO-GFP)^ fluorescence was imaged using a Nikon AXR microscope equipped with an NSPARC detector and a 60× oil-immersion objective (NA 1.42). Images were acquired using Nyquist and identical z-stack acquisition parameters across samples. Image stacks were deconvolved using Nikon imaging software and imported into Imaris version 10.2. For statistical analysis N numbers (biological replicates), statistical tests and p values for each count are indicated in the figure legends. Graphs and statistics were generated using GraphPad Prism Software 9 (GraphPad Inc., La Jolla, CA)

## Supporting information

SI Stats Table 1

SI Stats Table 2

SI Stats Table 3

## Data Availability

Raw and processed RNA-seq data have been deposited in the National Center for Biotechnology Information Gene Expression Omnibus (NCBI GEO), accessible through (GSE352267).

## Acknowledgments

This work was supported by NIH grant R01-EY035888 and a grant from the Retina Research Foundation to JMG, a William Shive Career Advancement Fellowship to RR, and NIH grant P20-GM144230 to MAR. Sequencing data was generated by the UTHealth Houston Cancer Genomics Core, supported by the Cancer Prevention and Research Institute of Texas (CPRIT RP240610). We’re grateful to the CBRS Microscopy and Flow Cytometry Core at UT Austin (RRID: SCR_021756) and particularly Michelle Mikesh for help with sample preparation, to Will Ramirez for zebrafish care, Yasaman Salmanpour for help with figure preparation, Fred Springsteen and members of the Gross lab for helpful suggestions on the project and manuscript.

## Notes

### Competing Interest Statement

The authors have declared no competing interest.

## References

1. Hoang, T., et al., Gene regulatory networks controlling vertebrate retinal regeneration. Science, 2020. 370(6519).

2. Jorstad, N.L., et al., Stimulation of functional neuronal regeneration from Müller glia in adult mice. Nature, 2017. 548(7665): p. 103–107.

3. Fausett, B.V., J.D. Gumerson, and D. Goldman, The proneural basic helix-loop-helix gene ascl1a is required for retina regeneration. J Neurosci, 2008. 28(5): p. 1109–17.

4. Khacho, M., et al., Mitochondrial Dynamics Impacts Stem Cell Identity and Fate Decisions by Regulating a Nuclear Transcriptional Program. Cell Stem Cell, 2016. 19(2): p. 232–247.

5. Fukuda, R., et al., Stimulation of glycolysis promotes cardiomyocyte proliferation after injury in adult zebrafish. EMBO Rep, 2020. 21(8): p. e49752.

6. Hutto, R.A., et al., Cone photoreceptors transfer damaged mitochondria to Müller glia. Cell Rep, 2023. 42(2): p. 112115.

7. Wogenstein, G.M., et al., Rod photoreceptors have a dual dependency on both aerobic glycolysis and OXPHOS and diverge metabolically from other retinal neurons. Sci Adv, 2026. 12(41): p. eaeg8687.

8. Beckers, A., et al., Optic nerve injury-induced regeneration in the adult zebrafish is accompanied by spatiotemporal changes in mitochondrial dynamics. Neural Regen Res, 2023. 18(1): p. 219–225.

9. Zhou, B., et al., Facilitation of axon regeneration by enhancing mitochondrial transport and rescuing energy deficits. J Cell Biol, 2016. 214(1): p. 103–19.

10. Yoshimatsu, T., et al., Presynaptic partner selection during retinal circuit reassembly varies with timing of neuronal regeneration in vivo. Nat Commun, 2016. 7: p. 10590.

11. Curado, S., et al., Conditional targeted cell ablation in zebrafish: a new tool for regeneration studies. Dev Dyn, 2007. 236(4): p. 1025–35.

12. Lee, M.S., J. Wan, and D. Goldman, Tgfb3 collaborates with PP2A and notch signaling pathways to inhibit retina regeneration. Elife, 2020. 9.

13. Krylov, A., et al., Heterogeneity in quiescent Müller glia in the uninjured zebrafish retina drive differential responses following photoreceptor ablation. Front Mol Neurosci, 2023. 16: p. 1087136.

14. Kanow, M.A., et al., Biochemical adaptations of the retina and retinal pigment epithelium support a metabolic ecosystem in the vertebrate eye. Elife, 2017. 6.

15. Weiss, E.R., et al., Broad spectrum metabolomics for detection of abnormal metabolic pathways in a mouse model for retinitis pigmentosa. Exp Eye Res, 2019. 184: p. 135–145.

16. Perez-Estrada, J.R., et al., Metabolic states influence chicken retinal pigment epithelium cell fate decisions. Development, 2024. 151(15).

17. Fausett, B.V. and D. Goldman, A role for alpha1 tubulin-expressing Müller glia in regeneration of the injured zebrafish retina. J Neurosci, 2006. 26(23): p. 6303–13.

18. Ramachandran, R., B.V. Fausett, and D. Goldman, Ascl1a regulates Müller glia dedifferentiation and retinal regeneration through a Lin-28-dependent, let-7 microRNA signalling pathway. Nat Cell Biol, 2010. 12(11): p. 1101–7.

19. Brew, H., et al., Endfeet of retinal glial cells have higher densities of ion channels that mediate K+ buffering. Nature, 1986. 324(6096): p. 466–8.

20. Dominguez-Romero, M.E., et al., Spinal cord injury induces mitochondrial dysfunction and metabolic reprogramming in non-regenerative Xenopus laevis. iScience, 2026. 29(7): p. 116636.

21. Anusha, P., et al., Aging compromises Zebrafish caudal fin regeneration by disrupting Regenerative gene networks and Cellular metabolism. bioRxiv, 2026: p. 2026.03.24.713633.

22. Cartoni, R., et al., The Mammalian-Specific Protein Armcx1 Regulates Mitochondrial Transport during Axon Regeneration. Neuron, 2016. 92(6): p. 1294–1307.

23. Lang, A.E., et al., Retrograde mitochondrial transport is required for mitochondrial biogenesis in zebrafish neurons. Nat Commun, 2026. 17(1).

24. Pan, X., et al., Mitochondrial dynamics govern whole-body regeneration through stem cell pluripotency and mitonuclear balance. Nat Commun, 2024. 15(1): p. 10681.

25. Zhao, Y., et al., Emerging roles of mitochondria in animal regeneration. Cell Regen, 2023. 12(1): p. 14.

26. Lin, Y.F., J. Sam, and T. Evans, Sirt1 promotes tissue regeneration in zebrafish through regulating the mitochondrial unfolded protein response. iScience, 2021. 24(10): p. 103118.

27. Hanna, J., et al., Glycolytic flux controls retinal progenitor cell differentiation via regulating Wnt signaling. Elife, 2025. 13.

28. Li, J., et al., Intron targeting-mediated and endogenous gene integrity-maintaining knockin in zebrafish using the CRISPR/Cas9 system. Cell Res, 2015. 25(5): p. 634–7.

