## Supplementary material for "Spatially patterned mitochondrial remodeling and metabolic changes during Müller glia-dependent retinal regeneration in zebrafish": SI Table 1

### Metric Interpretation Rules - 3D Mitochondrial Morphometry

| Metric | What it measures | How to interpret | Rule in tables | Conclusion when present |
|---|---|---|---|---|
| **Volume**<br>3D size | Total segmented mitochondrial 3D size | Reflects mitochondrial object size and content; not sufficient alone to distinguish expansion from elongation | Always interpreted with Area or Bounding Box | **Volume ↑ + Area ↑ = Expansion**<br>Volume ↑ + Bounding Box ↑ = Elongation |
| **Area**<br>Surface | Surface area of segmented mitochondria | Supports a size-increase interpretation; not valid in isolation without Volume | Interpreted only alongside Volume | **Area ↑ + Volume ↑ = Expansion** |
| **Bounding Box**<br>Length C - longest axis | Longest axis of the segmented object | Reflects spatial extension, not true connectivity. A long bounding box with high Volume supports elongation | Interpreted only alongside Volume | **Bounding Box ↑ + Volume ↑ = Elongation / extension** |
| **Sphericity**<br>Shape | Shape descriptor: roundness versus irregularity and elongation (0 = rod-like; 1 = perfect sphere) | Reflects morphology independently of size and captures shape remodeling even when Volume is unchanged | Interpreted independently or alongside other metrics | **Sphericity change = Morphology change independent of size** |
