## Supplementary material for "Spatially patterned mitochondrial remodeling and metabolic changes during Müller glia-dependent retinal regeneration in zebrafish": SI Materials and Methods

### Generation of gfap$^{Tg(MITO\text{-}GFP)}$

A mitochondrial targeting sequence derived from cytochrome *c* oxidase subunit VIII (COX8) fused to GFP (COX8-GFP) was directionally cloned into the vector using BamHI and AgeI restriction sites. Correct insert orientation and sequence integrity were confirmed by Sanger sequencing.

For embryo microinjection, CRISPR reagents were assembled as ribonucleoprotein (RNP) complexes immediately before use. Lyophilized sgRNA was resuspended in duplex buffer to a stock concentration of 200μM. Recombinant Cas9 protein (Integrated DNA Technologies) was diluted to 57μM in duplex buffer. Equal volumes (1μL each) of sgRNA and Cas9 were combined and incubated at 37°C for 5 minutes to form RNP complexes at a final concentration of 28.5μM.

The RNP complex was subsequently mixed with the donor plasmid to achieve a final plasmid concentration of 15 ng/μL. One nanoliter of the injection mixture was delivered into the yolk of one-cell-stage wild-type embryos before cell inflation, corresponding to approximately 4,700pg Cas9 protein, 1,000pg sgRNA, and 15pg donor plasmid per embryo.

Injected embryos were raised to adulthood, and F0 founders were screened for GFP fluorescence. Positive founders were outcrossed with wild-type AB fish to establish stable F1 transgenic lines, which were identified by GFP fluorescence and maintained by outcrossing to AB in subsequent experiments.

### Photoreceptor Ablation and Dichloroacetate (DCA) Treatment

Immediately following the 16h MTZ exposure, fish were rinsed three times with fresh system water and subsequently incubated in fresh system water for 10 min to facilitate removal of residual MTZ. Fish were then transferred to fresh system water containing 70mM DCA and maintained under standard husbandry conditions until tissue collection. Fresh DCA solution was prepared and replaced every 24h throughout the experimental period. Control animals were maintained under identical husbandry conditions in system water without MTZ or DCA. Retinae were collected at experimentally-defined time points following the initiation of MTZ treatment for immunohistochemistry, fluorescence-activated cell sorting (FACS), bulk RNA sequencing, metabolomic profiling, and super-resolution imaging

**Bulk RNA Sequencing of FACS-Sorted Müller Glia**

Retina enzymatically dissociated in oxygenated Ames' medium/PBS containing papain (20 U/mL), DNase I (80 U/mL), and L-cysteine (1.5mM) for 40 minutes at 28°C. Following enzymatic digestion, retinal tissue was gently triturated using a fire-polished glass Pasteur pipette to obtain a single-cell suspension. Enzymatic activity was terminated by addition of ovomucoid papain inhibitor supplemented with bovine serum albumin (BSA). The resulting cell suspension was passed through a 30-µm cell strainer, centrifuged at 250 × g for 8 minutes at 4°C, and the cell pellet was resuspended in Ames' medium containing 0.4% BSA and 1 µL DNase I.

Retinal cells isolated from age-matched wild-type (non-transgenic) zebrafish were used to establish background fluorescence and define GFP-positive sorting gates. Cell viability was assessed using DRAQ7 (Biostatus). Approximately 1,000 viable GFP-positive Müller glia were sorted directly into SMART-Seq HT lysis buffer (Takara Bio).

Full-length cDNA was synthesized using the SMART-Seq HT kit (Takara Bio) according to the manufacturer's instructions. Reverse transcription was performed using oligo(dT) priming and SMART template-switching technology to generate full-length cDNA from polyadenylated RNA. Sequencing libraries were prepared at the UTHealth Houston Genomics Core using the Illumina Nextera XT DNA Library Preparation Kit (Illumina). Paired-end sequencing (2 × 100 bp; 200-cycle chemistry) was performed on an Illumina NextSeq 2000 platform, targeting approximately 20 million reads per sample.

Raw paired-end FASTQ files were processed on the Center for Computational Biology and Bioinformatics (CCBB) high-performance computing cluster at the University of Texas at Austin. Adapter sequences and low-quality bases were removed using fastp. Filtered reads were aligned to the *Danio rerio* reference genome (GRCz11; Ensembl release 107) using HISAT2 (v2.2.1) with default parameters. SAM alignment files were converted to sorted and indexed BAM files using SAMtools (v1.12). Gene-level read counts were generated using featureCounts (Subread v2.0.6) in paired-end mode with the corresponding Ensembl release 107 gene annotation (GTF).

Raw count matrices were analyzed using the integrated Differential Expression and Pathway analysis (iDEP) platform (v0.96; Ge et al., 2018). Genes with low expression were excluded using the default RNA-seq filtering criteria implemented in iDEP. Count data were normalized using the DESeq2 normalization method and variance-stabilized using the variance stabilizing transformation (VST). Differential gene expression analysis was performed using the DESeq2 statistical framework implemented within iDEP. Genes with a Benjamini-Hochberg adjusted *P* value < 0.05 were considered significantly differentially expressed.

Differential expression analyses were performed by comparing each MTZ-treated time point (12, 24, 36, 48, and 60hpi) with uninjured controls, as well as each MTZ plus dichloroacetate (DCA) treatment group with its corresponding MTZ-treated control.

**Immunohistochemistry**

Eyes were dissected in ice-cold PBS supplemented with Pierce Phosphatase Inhibitor Mini Tablets (Thermo Fisher Scientific, Cat. No. A32957) and fixed in 4% PFA containing phosphatase inhibitor for 5h on ice. Following fixation, eyes were cryoprotected by sequential incubation in 25% sucrose overnight and 35% sucrose overnight at 4°C. Phosphatase inhibitors were maintained throughout the cryoprotection procedure for phospho-PDHα1 samples. Tissues were embedded in optimal cutting temperature (OCT) compound, cryosectioned at a thickness of 10 μm, mounted onto Superfrost Plus microscope slides, and stored at −20°C until use.

Cryosections were air-dried overnight at 4°C before antigen retrieval. Slides were immersed in preheated 10mM sodium citrate buffer (pH 6.0) and incubated at 90°C for 10min using a Coplin staining jar placed in a 500-mL beaker containing approximately 400 mL of buffer. Sections were allowed to cool in the same buffer for 20min at room temperature before excess buffer was removed and tissue sections were outlined using a hydrophobic barrier pen. Sections were rehydrated in PBS for 5 min and permeabilized with 0.5% Triton X-100 in PBS for 30 min at room temperature. Samples were blocked with 5% normal goat serum (NGS) in PBS for 1.5–2h at room temperature before incubation with mouse anti-PCNA primary antibody (clone PC10; Calbiochem/MilliporeSigma, Cat. No. NA03) diluted 1:100 in blocking solution overnight at 4°C. Following primary antibody incubation, sections were washed in PBS and incubated with goat anti-mouse IgG (H+L) Alexa Fluor 647 secondary antibody (Invitrogen/Thermo Fisher Scientific, Cat. No. A-21235) diluted 1:300 for 2h at room temperature. Sections were mounted using Vectashield Antifade Mounting Medium containing DAPI (Vector Laboratories). GFP fluorescence from the Tg(*tuba1a*:GFP) transgene was imaged directly without immunolabeling.

To preserve phospho-epitopes, all procedures performed on the first day of staining were carried out at 4°C. Cryosections were washed three times for 5min each in cold PBS containing 0.1% Tween-20 and permeabilized in 0.5% Triton X-100 in PBS for 30min on ice. Sections were blocked in 3% bovine serum albumin (BSA) prepared in PBS containing 0.1% Tween-20 for 1h at room temperature. Samples were incubated overnight at 4°C with rabbit monoclonal anti-phospho-pyruvate dehydrogenase E1α (Ser293) antibody (clone E4V9L; Cell Signaling Technology, Cat. No. 37115) diluted 1:200 in blocking buffer. On the second day, sections were washed three times for 10min each in PBS containing 0.1% Tween-20 and incubated with goat anti-rabbit IgG Alexa Fluor 647 secondary antibody (Invitrogen/Thermo Fisher Scientific, Cat. No. A-21244) diluted 1:200 for 2h at room temperature while protected from light. Sections were washed three additional times for 10min each before mounting with ProLong Gold Antifade Mountant containing DAPI (Thermo Fisher Scientific).

**Sample Preparation and Immunolabeling for NSPARC Super-Resolution Imaging**

Dissected retinae were fixed in 4% paraformaldehyde (PFA) prepared in Ames' medium for 25 min on ice. Following fixation, residual aldehydes were quenched by incubation in 100 mM glycine for 10 min at 4°C. Tissues were subsequently blocked and permeabilized in Ames' medium containing 10% normal goat serum (NGS) and 0.5% Triton X-100 for 3h at 4°C. Images were obtained using a 60x oil objective (NA = 1.42).

For NSPARC imaging, retinae were incubated with rabbit polyclonal anti-GFP antibody (Invitrogen/Thermo Fisher Scientific, Cat. No. A-11122; RRID: AB_221569) in blocking solution (5μg per retina) for 24h at 4°C with gentle agitation. The primary antibody solution was then replaced with freshly prepared antibody at the same concentration, and tissues were incubated for an additional 48h to maximize antibody penetration and labeling efficiency. Following primary antibody incubation, retinae were washed six times for 10min each in Ames' medium containing 2% NGS. Samples were then incubated overnight (12–16h) at 4°C with goat anti-rabbit Alexa Fluor 647 secondary antibody (Invitrogen/Thermo Fisher Scientific, Cat. No. A-21244) at 8 μg/mL in 2% NGS supplemented with 1× protease inhibitor cocktail. After secondary antibody labeling, tissues were washed six times for 5min each in 2% NGS prepared in Ames' medium.

**Resin Embedding and Sectioning**

After immunolabeling, all retinas were post-fixed in 2% PFA and 0.5% glutaraldehyde in PBS for 30 min at 4°C with gentle agitation. Samples were dehydrated through a graded ethanol series (50%, 70%, 90%, 100%, and 100%) for 15 min at each step while gently rotating at room temperature. Dehydrated tissues were infiltrated with Ultra Bed Low Viscosity Epoxy Resin (Electron Microscopy Sciences, Cat. No. 14310) using a stepwise resin:ethanol series consisting of 1:3 resin:ethanol for 2 h, 1:1 resin:ethanol for 2 h, and 3:1 resin:ethanol overnight (~16 h), all with gentle rotation at room temperature. Samples were subsequently incubated twice in 100% resin for 2 h each before being transferred to embedding molds and polymerized at 65°C for 24 h. For NSPARC imaging, 1-µm-thick semithin sections were cut through the central retina using a Leica UC7 ultramicrotome and mounted onto glass microscope slides.

**Whole eye cup Metabolomics**

Immediately following dissection, eye cup were snap-frozen in liquid nitrogen and stored at −80°C until shipment to Metabolon Inc. (Morrisville, NC, USA) for metabolomic analysis. To characterize metabolic changes associated with retinal regeneration, global untargeted metabolomic profiling was performed on eye cup collected from adult zebrafish (3–5 months of age) at six experimental time points: uninjured control and 12, 24, 36, 48, and 60 hours post-injury (n = 4 biological replicates per time point). Approximately 30 mg of retinal tissue was collected for each biological replicate. Immediately following dissection, eye cup were snap-frozen in liquid nitrogen and stored at −80°C until shipment to Metabolon Inc. (Morrisville, NC, USA) for metabolomic analysis. Global untargeted metabolomic profiling was performed by Metabolon Inc. using ultrahigh-performance liquid chromatography coupled with tandem mass spectrometry (UPLC-MS/MS). Samples were extracted and analyzed using a Waters ACQUITY UPLC system coupled to a Thermo Scientific Q Exactive high-resolution mass spectrometer equipped with a heated electrospray ionization (HESI-II) source and Orbitrap mass analyzer operated at a resolution of 35,000. To maximize metabolite coverage, four complementary chromatographic methods were employed: two reverse-phase C18 methods operated under acidic positive-ion conditions optimized for hydrophilic (Pos Early) and hydrophobic (Pos Late) compounds, one reverse-phase C18 method operated under basic negative-ion conditions, and one hydrophilic interaction liquid chromatography (HILIC) method operated under negative-ion conditions. Data were acquired across a mass-to-charge (m/z) range of 70–1000. Raw spectral data were processed using Metabolon's proprietary informatics pipeline. Metabolites were identified by comparison with Metabolon's reference library of more than 5,400 authenticated chemical standards using three independent criteria: retention index, accurate mass (±10 ppm), and MS/MS spectral matching. Metabolite abundance was quantified using peak area under the curve, followed by median block normalization to correct for inter-day instrument variation. Missing values were imputed using the minimum observed value for each metabolite. Statistical analyses were performed on log-transformed, median-scaled metabolite abundances using Welch's two-sample *t*-test. Metabolites with $P \leq 0.05$ were considered significantly altered, whereas metabolites with $0.05 < P < 0.10$ were considered to exhibit a statistical trend. False discovery rates were estimated by calculating *q*-values according to the method of Storey and Tibshirani (2003). Differential metabolite abundance was assessed by comparing each post-injury time point (12, 24, 36, 48, and 60 hours post-injury) with uninjured controls.

**Image acquisition, quantification**

Three-dimensional surfaces of mitochondrial fluorescence were generated using the Surfaces tool. Surface smoothing was enabled with a surface detail value of 0.100 µm. Segmentation was performed using background subtraction (local contrast), with the "diameter of largest sphere which fits into the object" parameter set to 0.350 µm and an intensity threshold adjusted. Surfaces were filtered by voxel number. The same segmentation settings were applied across all images. For three-dimensional mitochondrial segmentation, mitochondrial volume, surface area, sphericity, and longest-axis length were extracted separately for the ganglion cell layer, inner plexiform layer, inner nuclear layer, outer plexiform layer, and outer nuclear layer. Volume, surface area, and longest-axis length were divided by the area of the corresponding retinal layer, measured in Fiji for the same image. Sphericity was analyzed without area normalization. Phospho-PDHA1 immunofluorescence was imaged using a Nikon AXR microscope with a 60× oil-immersion objective (NA 1.42). The same number of optical sections was acquired for each sample, and maximum-intensity projections were generated in Fiji. Mean fluorescence intensity was measured within the retinal region of interest. Background correction was performed by subtracting the mean fluorescence intensity of a region lacking phospho-PDHA1 signal within the same image. PCNA and *tuba1a* fluorescence images were acquired using a Nikon AXR microscope with 40× (NA 1.3) and 20× (NA 0.8) objectives. Z-stack acquisition parameters were maintained across samples, and maximum-intensity projections were generated in Fiji. PCNA-positive and *tuba1a* cells were counted manually using images acquired with the 40× objective. Individual optical sections were also inspected to identify cells obscured in the projections.

**Manuscript preparation:**

All text in this manuscript was written by the authors. ChatGPT (GPT-6) was used in editing sections of the text to improve the clarity. The authors take full responsibility for the published content.
