## Supplementary material for "Spatially patterned mitochondrial remodeling and metabolic changes during Müller glia-dependent retinal regeneration in zebrafish": SI Figures

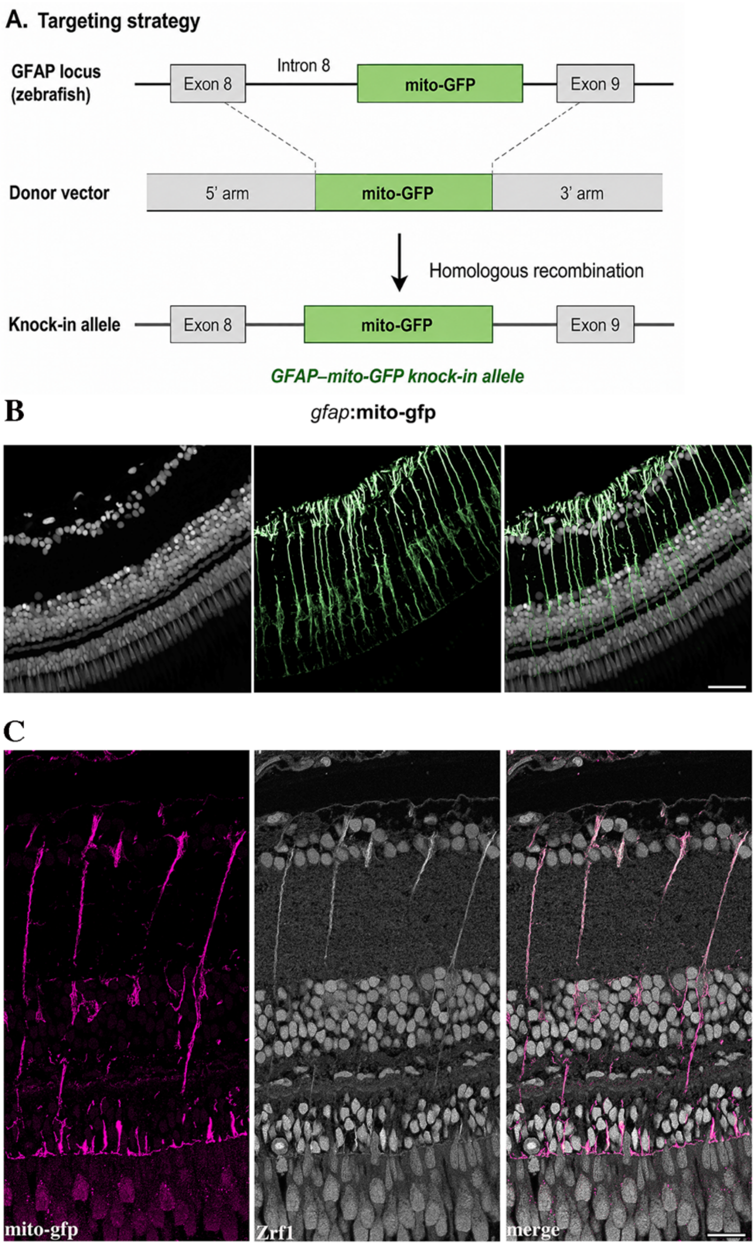


**Figure S1. Generation and characterization of a Müller glia-specific mitochondrial reporter line.**

Targeting strategy used to generate the *gfap*$^{Tg(MITO\text{-}GFP)}$ knock-in allele. A donor construct containing a mitochondrially targeted GFP flanked by 5′ and 3′ homology arms was integrated into intron 8 of the zebrafish *gfap* locus, between exons 8 and 9, by homologous recombination. (B) Representative 10-μm retinal section from adult *gfap*$^{Tg(MITO\text{-}GFP)}$ zebrafish showing nuclei in grayscale, mito-GFP fluorescence in green, and the merged image. Mito-GFP labels Müller glial compartments across the retinal layers. (C) Representative 2-µm resin-embedded retinal section showing colocalization of mito-GFP with the Müller glial marker Zrf1. Mito-GFP is shown in magenta, Zrf1 immunoreactivity in grayscale, and the merged image is shown at right. Scale bars = 20 µm (B) and 10 µm (C).

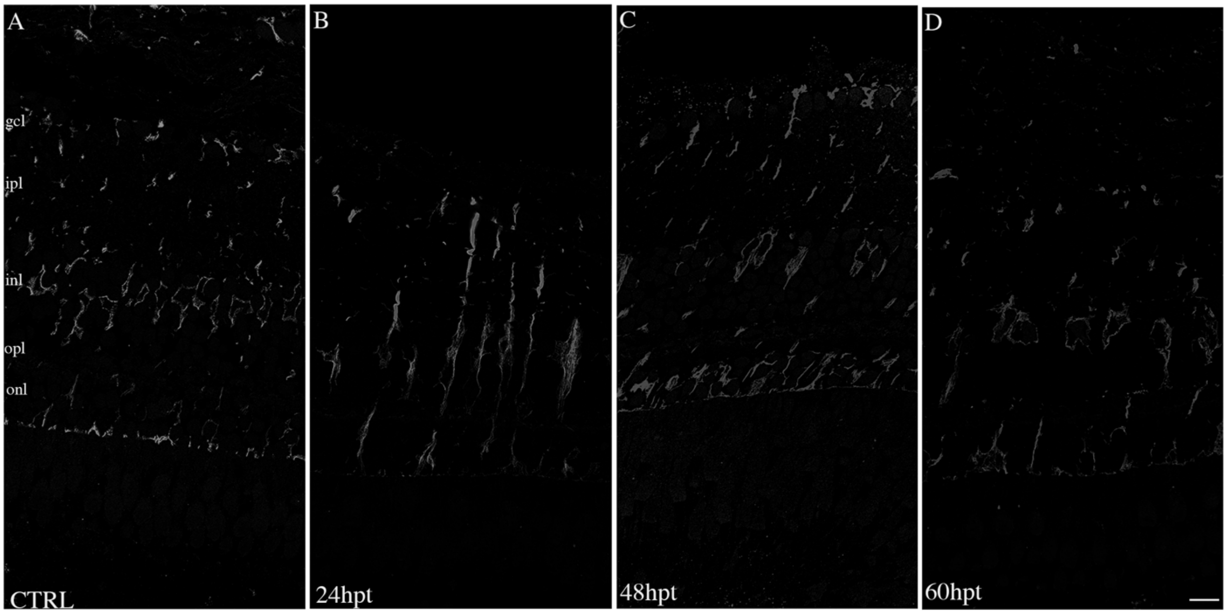


**Figure S2. Representative source images for analysis of Müller glial mitochondrial morphology after photoreceptor loss.**

Representative images of 1-μm resin-embedded retinal sections from adult *gfap*$^{\mathrm{Tg(MITO\text{-}GFP)}}$*;sws1:nfsB-mCherry* zebrafish. Sections were immunostained for GFP and imaged using an NSPARC-equipped microscope. (A) Untreated control retina. (B–D) Retinae collected at 24 (B), 48 (C), and 60hpt (D). Mito-GFP immunoreactivity is shown in gray. These source images were used for three-dimensional segmentation and morphometric analysis of mitochondrial objects in Imaris. Scale bar = 10 μm.

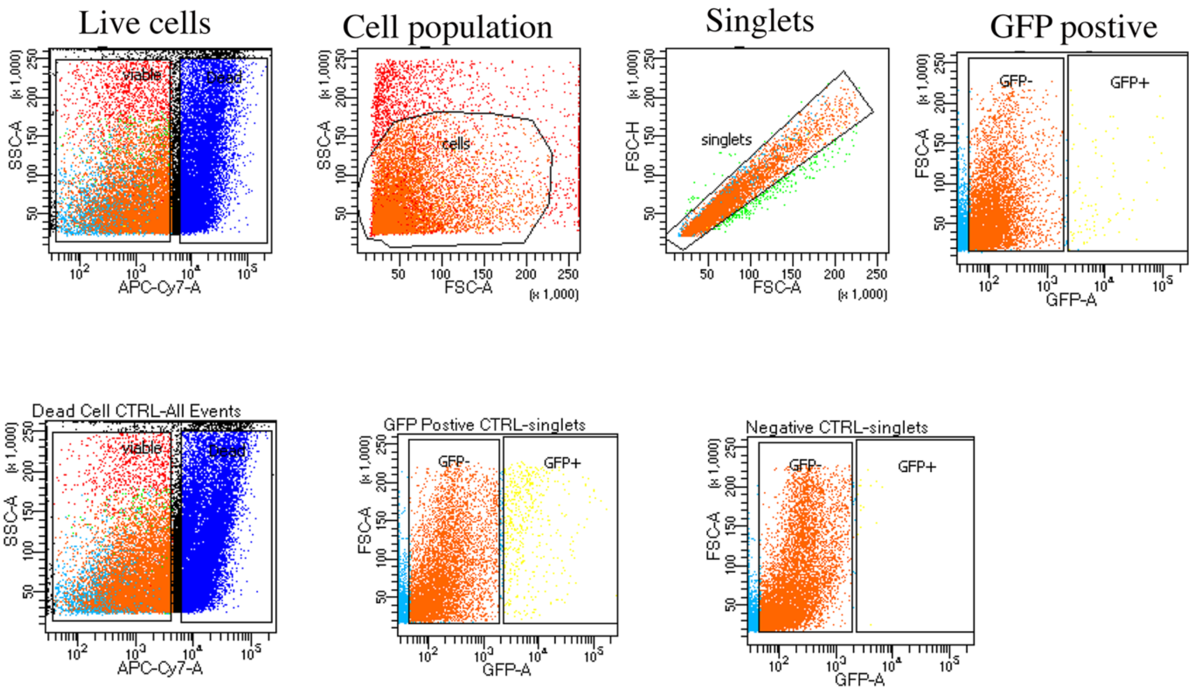


**Figure S3. Fluorescence-activated cell-sorting strategy for isolation of *gfap*:GFP Müller glia.**

Representative flow-cytometry plots showing the gating strategy used to isolate GFP-positive Müller glia from dissociated adult zebrafish retinae for RNA sequencing. Viable cells were using a viability dye detected in the APC-Cy7 channel and side-scatter area (SSC-A). The principal cell population was selected using forward-scatter area (FSC-A) and SSC-A, followed by exclusion of doublets and aggregates using FSC-A and forward-scatter height (FSC-H). GFP-positive cells were subsequently identified using GFP fluorescence intensity and FSC-A and collected for RNA isolation and sequencing. Lower plots show the dead-cell control, GFP-positive and GFP-negative control used to establish the gates. FSC, forward scatter; SSC, side scatter; FACS, fluorescence-activated cell sorting.

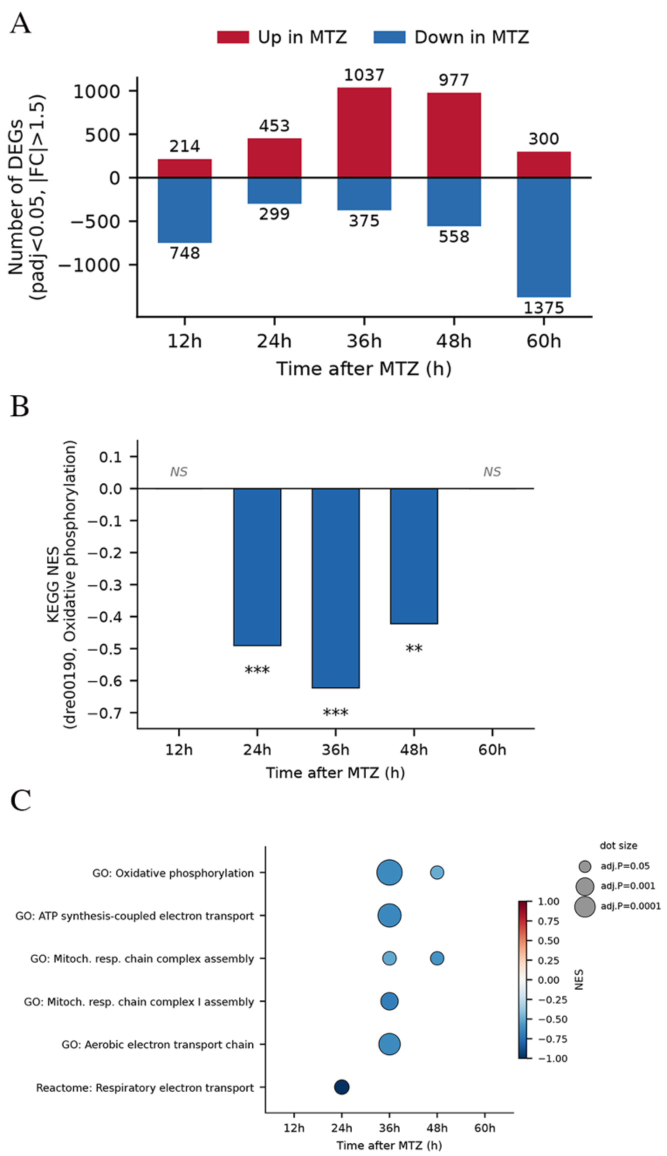


**Figure S4. Photoreceptor loss induces a staged Müller glial transcriptional response.**

(A) Numbers of differentially expressed genes in FACS-isolated Müller glia at 12, 24, 36, 48, and 60hpt relative to untreated controls. Differential expression was defined using an absolute fold-change > than 1.5 and a Benjamini–Hochberg-adjusted *P* value < 0.05. Red bars indicate increased expression, and blue bars indicate reduced expression. The numbers of upregulated and downregulated genes, respectively, were 214 and 748 at 12hpt, 453 and 299 at 24hpt, 1,037 and 375 at 36hpt, 977 and 558 at 48hpt, and 300 and 1,375 at 60hpt. (B) Gene-set enrichment analysis of the KEGG oxidative phosphorylation pathway (dre00190). Bars show normalized enrichment scores (NES) for MTZ-treated Müller glia relative to untreated controls; negative NES values indicate enrichment among genes reduced following injury. Oxidative phosphorylation was Significant negative enrichment was detected at 24, 36, and 48hpt. (C) Dot plot showing significantly enriched Gene Ontology and Reactome gene sets associated with oxidative phosphorylation, ATP synthesis-coupled electron transport, mitochondrial respiratory-chain assembly, and respiratory electron transport. Dot color indicates NES, with blue representing negative enrichment; and dot size reflects the adjusted *P* value. Enrichment significance was assessed using multiple-testing-adjusted *P* values. **P < 0.01 and *** *P* < 0.001; NS, not significant.

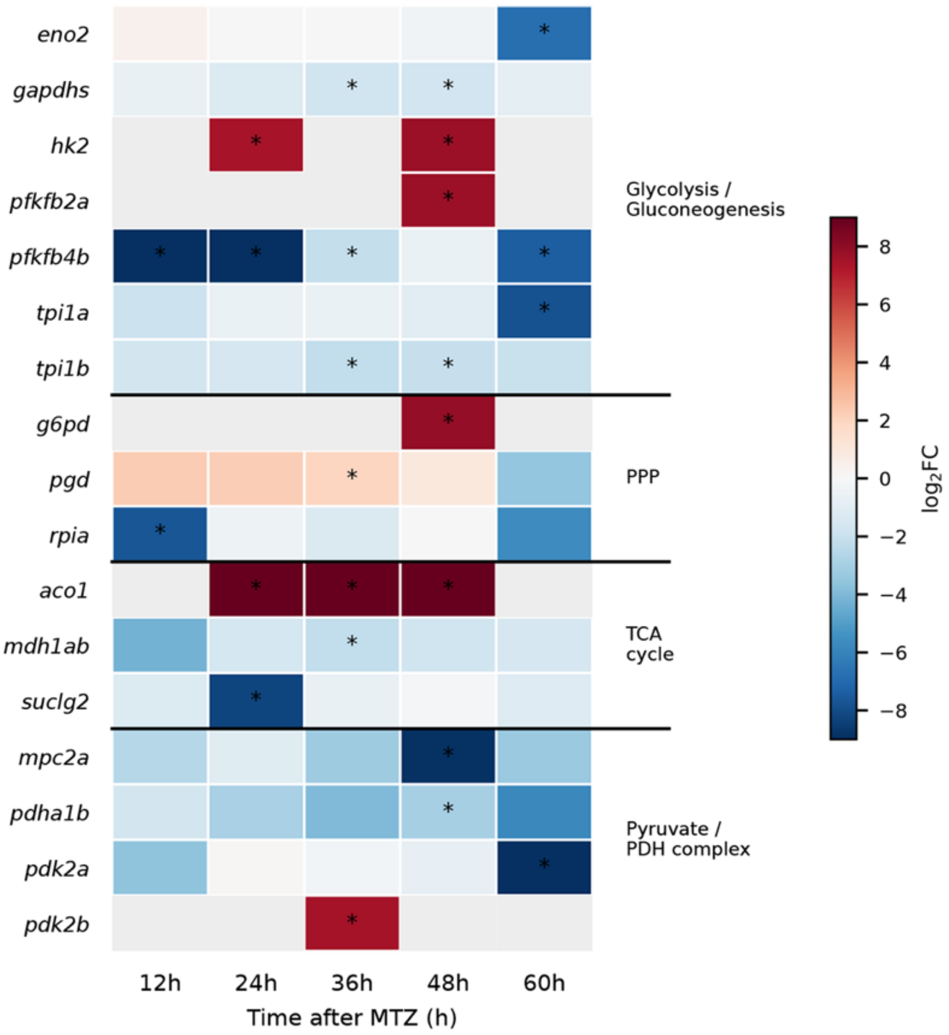


**Figure S5. Photoreceptor loss induces temporally heterogenous changes in Müller glia metabolic gene expression.**

Heatmap showing changes in expression of selected genes associated with glycolysis/gluconeogenesis, the pentose phosphate pathway (PPP), the tricarboxylic acid (TCA) cycle, and pyruvate metabolism/PDH regulation in FACS-isolated Müller glia at 12, 24, 36, 48, and 60hpt. Values represent $\log_2$ fold changes relative to uninjured controls; red denotes increased expression and blue denotes reduced expression. Asterisks indicate significant differential expression (Benjamini–Hochberg-adjusted $P < 0.05$).

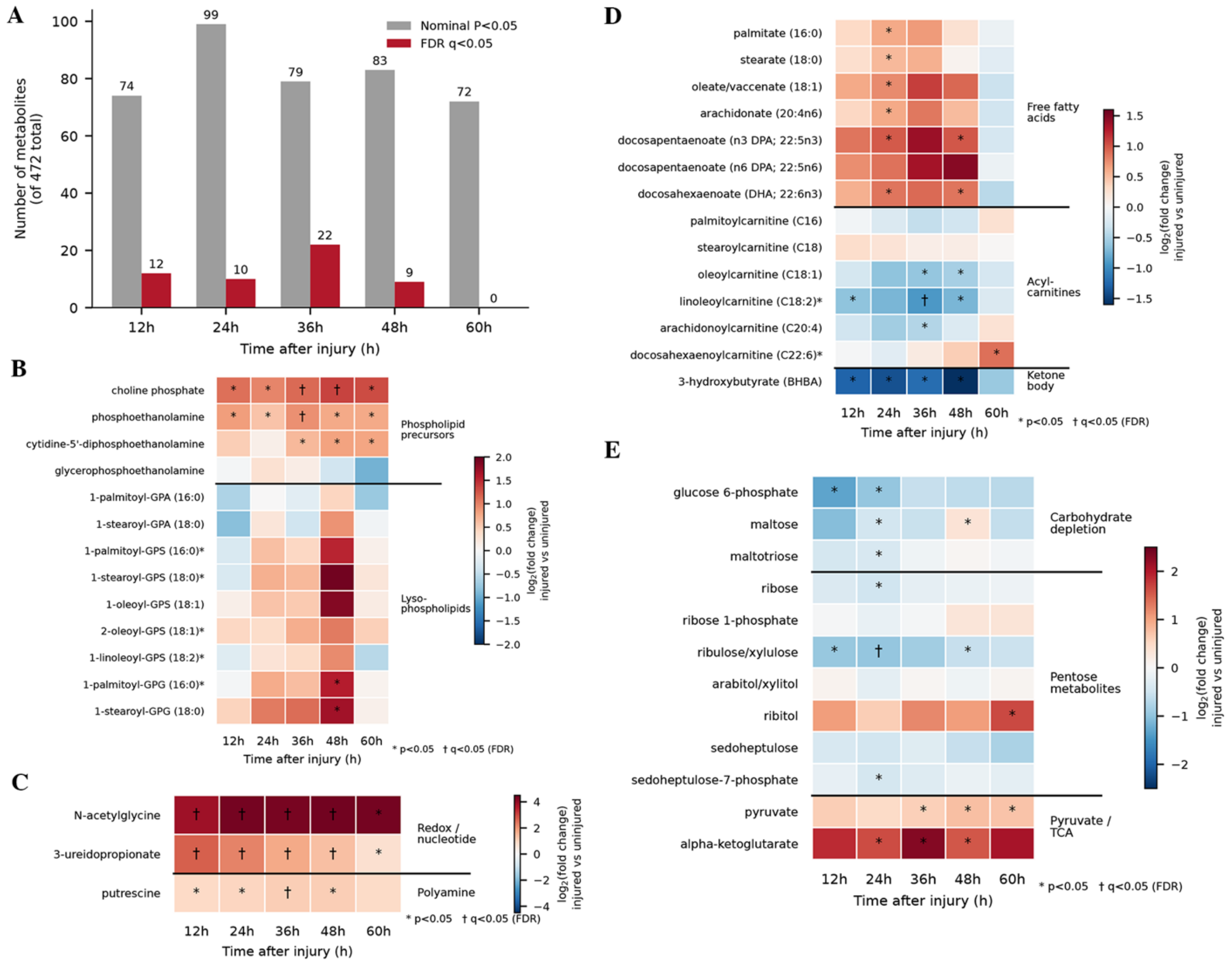


**Figure S6. Photoreceptor loss induces temporally patterned changes in the eye-cup metabolome.**

Untargeted metabolomic profiling was performed on whole eye cups collected at 12, 24, 36, 48, and 60hpt and compared with time-matched uninjured controls (n=4 replicates per condition). (A) Numbers of metabolites differing between injured and uninjured eye cups among the 472 detected metabolites. Gray bars indicate metabolites meeting the nominal significance threshold ($P < 0.05$), and red bars indicate significant after false-discovery-rate correction ($q < 0.05$). (B–E) Heatmaps showing changes in selected metabolites associated with (B) phospholipid precursors and lysophospholipids; (C) redox, nucleotide and polyamine metabolism; (D) free fatty acids, acylcarnitines, and ketone-body metabolism; and (E) carbohydrate, pentose, pyruvate, and TCA-cycle metabolism. Values represent $\log_2$ fold changes in injured relative to uninjured controls**.** Statistical comparisons were performed Welch's two-sample t test Asterisks indicate nominal $P <$

0.05; daggers indicate FDR $q < 0.05$. Exact $P$ and $q$ values and the complete metabolomics dataset are provided in SI table 4.

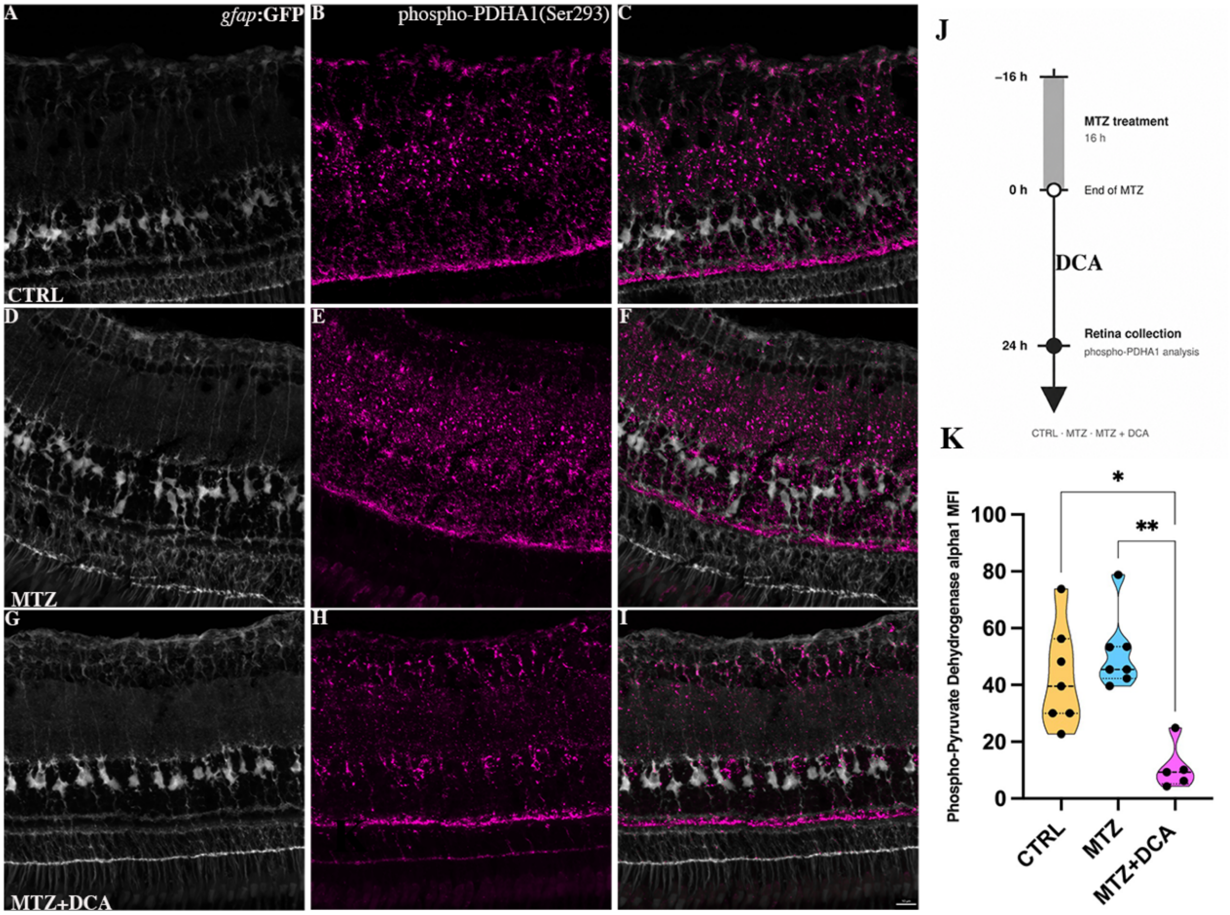


**Figure S7. DCA reduces retinal PDHA1 phosphorylation after photoreceptor injury.**

(A–I) Representative retinal sections from untreated control (A–C), MTZ-treated (D–F), and MTZ+DCA-treated (G–I) Tg(*gfap*:GFP)*;*Tg(*sws1*:nfsB-mCherry) zebrafish collected 16h after completion of MTZ exposure. GFP-labeled Müller glia are shown in grayscale (A,D,G), phospho-PDHA1 (Ser293) immunoreactivity in magenta (B,E,H), and in merged images in (C,F,I). (J) Experimental timeline. (K) Quantification of phospho-PDHA1 (Ser293) mean fluorescence intensity (MFI). Image quantification from three central sections per retina. Each point represents one biological sample: CTRL n=7; MTZ n=8; MTZ+DCA n=6. Violin plots show the median and quartiles. Statistical significance was assessed using one-way ANOVA followed by Tukey's multiple-comparisons test. $*P < 0.05$; $**P < 0.01$. Scale bar = 10 µm.

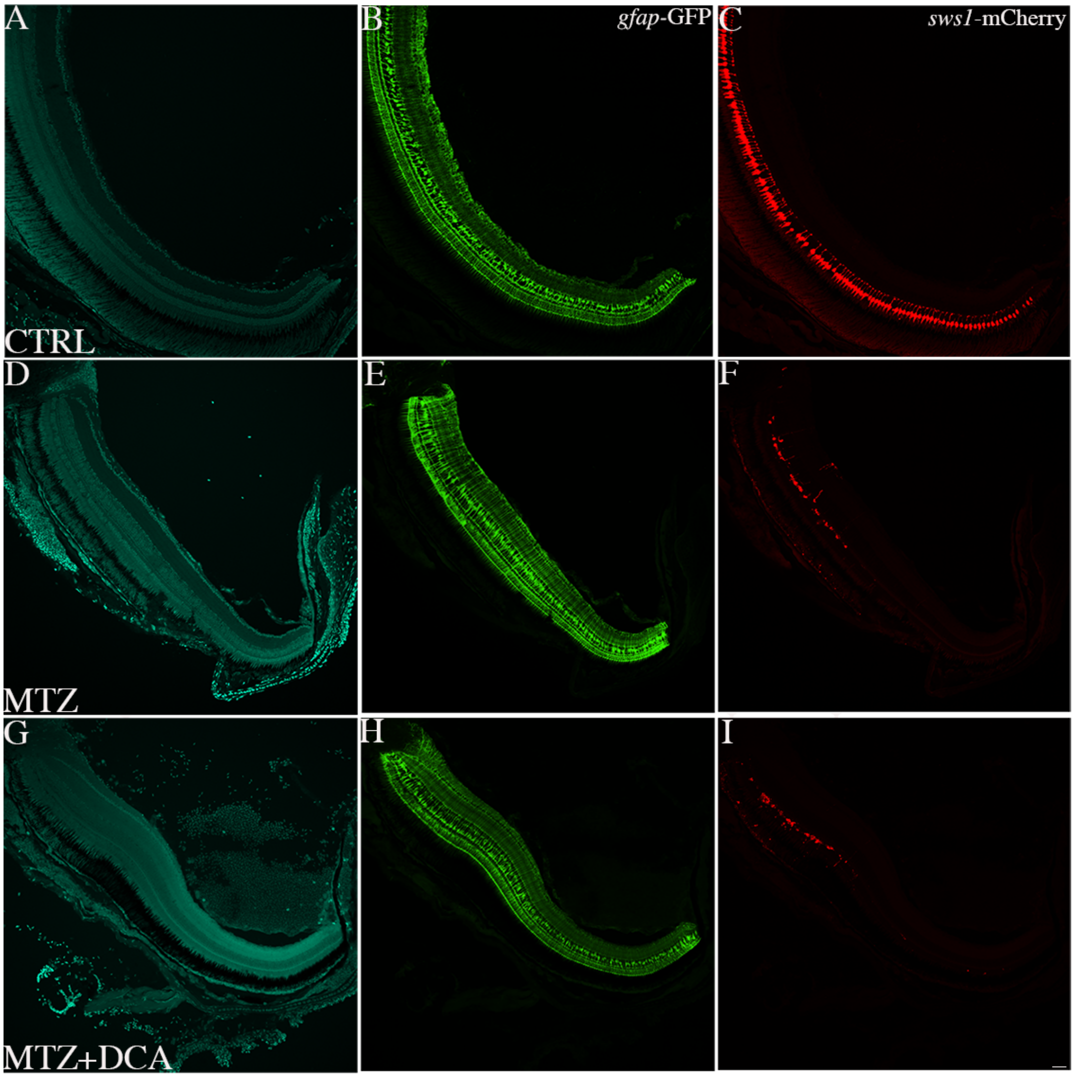


**Figure S8. UV cone loss following MTZ treatment in the presence or absence of DCA.**

Representative retinal sections from uninjured controls (A–C), MTZ-treated fish (D–F), and MTZ+DCA-treated fish (G–I), samples collected at 48 hours post MTZ treatment (hpt). Nuclear staining is shown in cyan (A, D, G), gfap-driven GFP in green (B, E, H), and *sws1*-driven mCherry labeling UV cones in red (C, F, I). Both MTZ and MTZ+DCA groups show a marked reduction in mCherry-positive UV cones relative to controls, with no obvious difference between MTZ vs MTZ+DCA. Scale bar = 50 μm.

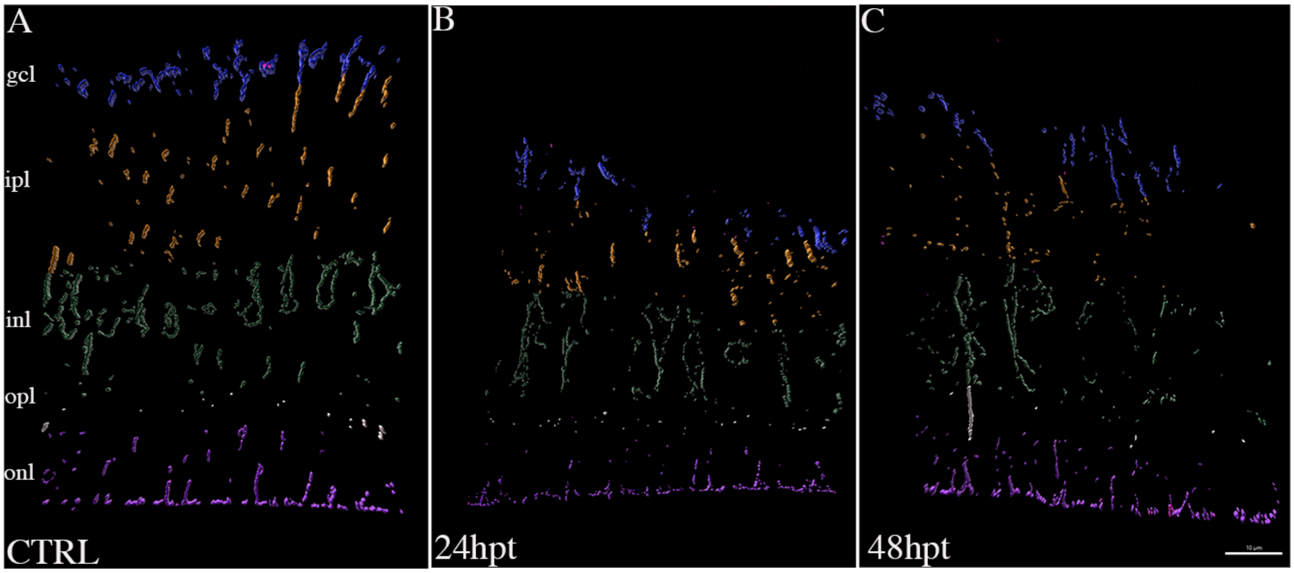


**Figure S9. Müller glial mitochondrial morphology following DCA treatment in the absence of UV cone ablation.**

Representative three-dimensional surface reconstructions of mito-GFP–labeled mitochondria in Müller glia from untreated controls (A) and at 24 (B) and 48 (C) hours posttreatment (hpt) with DCA. Mitochondrial surfaces are pseudocolored according to retinal compartment. Representative images show no obvious alteration in overall mitochondrial morphology following DCA treatment without UV cone ablation. Scale bar=10 µm.

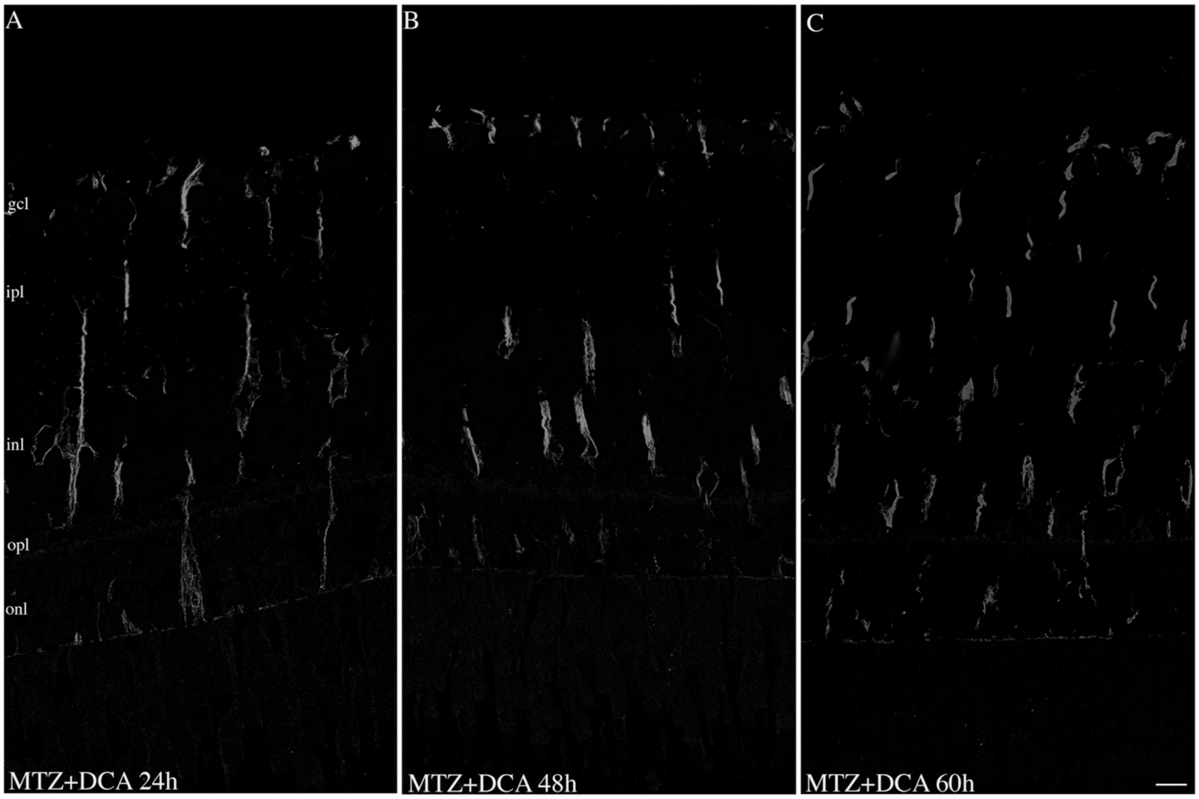


**Figure S10. Representative source images used for three-dimensional analysis of Müller glial mitochondria following DCA treatment.**

Representative images of 1-µm resin-embedded retinal sections from adult *gfap*$^{Tg(MITO-GFP)}$*;sws1:nfsB-mCherry* zebrafish. Sections were immunostained for GFP and imaged using an NSPARC-equipped microscope. (A–C) Retinae collected at 24 (A), 48 (B), and 60hpt (C) treated with DCA. Mito-GFP immunoreactivity is shown in gray. These source images were used for three-dimensional segmentation and morphometric analysis of mitochondrial objects in Imaris. Scale bar = 10 µm.

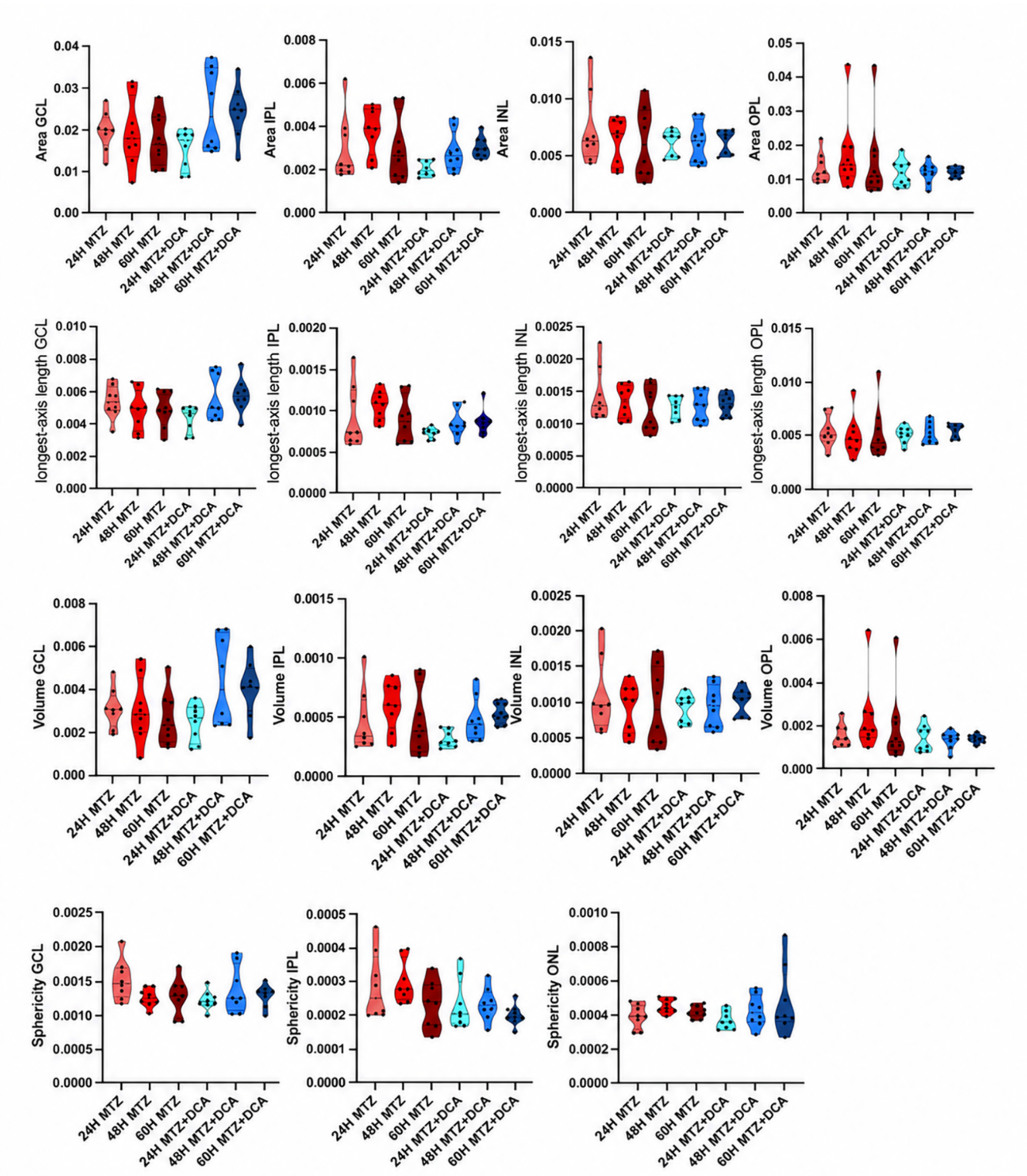


**Figure S11. Additional analyses of Müller glial mitochondrial morphology following DCA treatment after photoreceptor loss.**

Violin plots showing mitochondrial surface area, longest-axis length, and volume in the ganglion cell layer (GCL), inner plexiform layer (IPL), inner nuclear layer (INL), and outer plexiform layer (OPL), and mitochondrial sphericity in the GCL, IPL, and outer nuclear layer (ONL), at 24, 48, and 60 hours postinjury. MTZ-treated fish were compared with fish receiving MTZ+DCA. Black dots indicate individual measurements. These analyses complement the outer-retinal mitochondrial measurements presented in Figure 5.

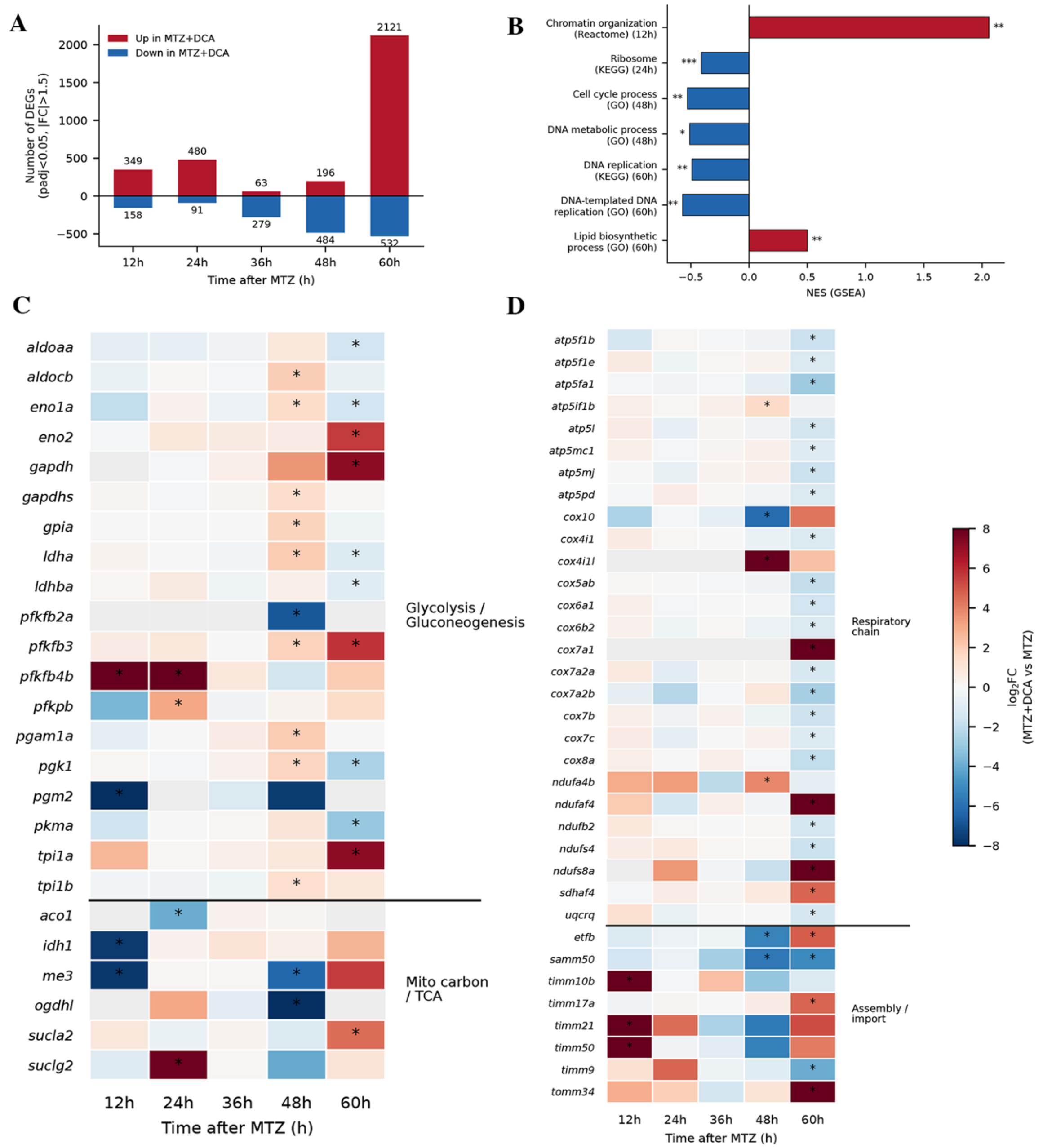


**Figure S12. DCA reshapes the Müller glia transcriptional response to photoreceptor loss.**

Transcriptomes of FACS-isolated Müller glia from MTZ+DCA-treated retinas were compared with those from time-matched MTZ-treated retinas at 12, 24, 36, 48, and 60hpt. (A) Numbers of differentially expressed genes at each time point using an adjusted $P$ value < 0.05 and an absolute fold change > 1.5 in either direction. Red bars indicate genes increased and blue bars indicate reduced genes expression with DCA treatment. (B) Gene-set enrichment analysis showing selected pathways significantly altered by DCA. Positive normalized enrichment scores (NES; red) indicate enrichment in MTZ+DCA-treated Müller glia; negative NES values (blue) indicate enrichment among genes with reduced expression relative to MTZ-treated Müller glia. The time point and annotation database are indicated for each pathway. (C,D) Heatmaps showing DCA-induced changes in genes associated with (C) glycolysis/gluconeogenesis and mitochondrial carbon metabolism/TCA-cycle function and (D) respiratory-chain components and mitochondrial protein-complex assembly/import. Values represent $\log_2$ fold changes in MTZ+DCA-treated samples relative to MTZ-treated samples; red denotes increased expression and blue denotes reduced expression. Asterisks in C,D indicate an adjusted $P$ value < 0.05. For enrichment analyses in B, adjusted $*P < 0.05$, $**\ P < 0.01$, and $***P < 0.001$, respectively.
